# Differential expression of tension-sensitive *HOX* genes in fibroblasts is associated with different scar types

**DOI:** 10.1101/2023.07.02.547450

**Authors:** Minwoo Kang, Ung Hyun Ko, Eun-Jung Oh, Hyun Mi Kim, Ho Yun Chung, Jennifer H. Shin

## Abstract

A scar is considered a natural consequence of the wound-healing process. However, the mechanism by which scars form remains unclear. Here, we suggest a new mechanism of wound healing and scar formation that involves the mechanosensitive regulation of *HOX* genes. RNA-sequencing of fibroblasts from different types of scars revealed differential *HOX* gene expression. Computational simulations predicted injury-induced tension loss in the skin, and in vitro experiments revealed a negative correlation between tension and fibroblast proliferation. Remarkably, exogenous tensile stress in fibroblasts has been shown to alter *HOX* gene expression levels in different scar types. Overall, we propose a model for normal wound healing and scar formation and show that successful wound healing requires tensional homeostasis in the skin tissue, which is regulated by tension-sensitive *HOX* genes.

## Introduction

The wound-healing process is a complex phenomenon that constantly occurs in living organisms. This process can be viewed as a part of tissue homeostasis, in which newly produced tissue can replace and restore damaged tissue. At the end of the process, the healing tissue can form nonfunctioning fibrotic regions, referred to as scars. Scar tissues are composed mainly of aligned collagen fibers, unlike the randomly weaved structure in normal tissue, and the alignment in the collagen network is generated by activated fibroblasts^1–3^. Among the many types of cells involved in the wound healing process, fibroblasts are the key players that drive the regeneration of damaged connective tissue. Wound healing consists of four interconnected phases: hemostasis, inflammation, proliferation, and tissue remodeling. During the initial inflammatory response mediated by white blood cells, fibroblasts migrate toward the wound site via a chemotactic response to inflammatory cytokines. Within a couple of days, fibroblasts at the wound site enter the proliferative phase^4^, followed by the tissue remodeling phase, in which fibroblasts reconstruct the extracellular matrix (ECM) by secreting matrix proteins^5^. During this process, some fibroblasts differentiate into myofibroblasts, an activated form of fibroblasts, to enhance the secretion of cytokines and ECM proteins while inducing tissue contraction^6^.

As a wound heals, some people develop abnormal scarring in the form of hypertrophic scars or keloids. These scars have an unappealing appearance that may affect patients physically and psychologically. Interestingly, hypertrophic scars and keloids share discolored skin extrusions of excessive collagen. However, these scars feature different degrees of severity and are traditionally classified as two different scar types. Scars that do not grow beyond the original wound boundaries and feature a thickened and raised morphology are defined as hypertrophic scars, whereas keloids are scars that spread into the surrounding wound edges with a similar but more aggressive morphology^7,8^.

In many cases, however, hypertrophic scars and keloids have similar growth and histological characteristics. The majority of researchers believe that keloids are a distinct clinical entity and a systemic disorder, as opposed to a scar or a severe hypertrophic scar. Since hypertrophic scars and keloid formation are influenced by diverse factors, including systemic, genetic, local, and lifestyle risk factors, understanding the factors that drive the formation and progression of these scars is essential for effective prevention and treatment ^9,10^. Meanwhile, another opinion holds that in histopathology, keloids and hypertrophic scars are manifestations of the same inflammatory fibroproliferative condition and vary in intensity and duration of inflammation ^11–13^. There are empirical clues concerning why such abnormal scars are formed, including genetic and mechanical effects. Interestingly, keloids have been shown to occur in genetically susceptible individuals^14,15^. For example, keloid development is more frequently observed in dark-skinned individuals^16^. Regarding the incidence of keloids after the caesarian section, African American and Asian individuals showed significantly increased keloid formation compared to Caucasian individuals^17^. Based on empirical evidence, it has been suggested that increased pigmentation could be a factor in keloid development^18,19^. However, African individuals with albinism showed no significant difference from those with normal pigmentation in terms of the prevalence rate of keloid formation, suggesting that increased pigmentation may not be the fundamental cause of keloid formation^20^. On the other hand, it is recognized that mechanical forces considerably influence the biological processes of wound healing and scar formation. Mechanotransduction is the process by which cells convert mechanical forces into biochemical signals^21,22^. Inadequate mechanotransduction has been linked to pathological wound healing, such as over-healing (fibrosis and scar formation) and under-healing (chronic wounds)^21,23^. Specifically, mechanical tension has been considered the leading cause of both hypertrophic scars and keloid formation based on the observation that most abnormal scars form in the skin under relatively high tension in a site-specific manner^24,25^. However, based on the development of keloids on the earlobes, where the tension is very low, this hypothesis only partially explains the influence of high tension on hypertrophic scar formation. In short, no clear scientific evidence exists concerning the genetic and mechanical mechanisms of scar formation.

The current study primarily centers on fibroblasts and their significant role in the excessive production of extracellular matrix (ECM), ultimately leading to the formation of scars. By examining both transcriptomic and mechanobiological aspects, we offer a novel perspective on the wound-healing process, providing further understanding of the formation of abnormal scars. We first isolated fibroblasts from patients and compared the differences in their transcriptomes. To our surprise, transcriptome profiling identified extraordinarily high expression levels of *HOX* genes in fibroblasts originating from hypertrophic scars compared to fibroblasts from either normal scars or keloids. Computational simulation suggested that injury-induced alterations in the tensional state occurred at the wound site. Thus, we hypothesized that the loss of tension could serve as the signal promoting fibroblast proliferation, thereby initiating the wound healing process. In vitro tensile stimulation experiments demonstrated that mechanical tension could suppress proliferation in both normal skin and hypertrophic scars but not in keloids. Furthermore, we found a positive correlation between tensile stimulation and *HOX* gene expression in normal fibroblasts. To our surprise, fibroblasts from hypertrophic scars and keloids exhibited entirely dissimilar responses to mechanical tension, and only hypertrophic scar fibroblasts featured a mechano-response similar to that of normal fibroblasts.

We integrate these key findings and propose a novel model to explain normal wound healing and scar formation. Our model holds that successful wound healing requires homeostatic maintenance of the intrinsic tensile stress (i.e., tensional homeostasis) in the skin tissue via the regulation of tension-sensitive *HOX* genes. With the proposed model, we can not only demonstrate how abnormal scars develop but also distinguish hypertrophic scars from keloids. Furthermore, the model can be used to develop strategic treatment approaches to effectively reduce scar formation.

## Results

### Fibroblasts from hypertrophic scars and keloids are distinguishable by morphological features but not by total mRNA expression

Normal skin, hypertrophic scar, and keloid scar tissues were acquired from three patients for each type. Then, fibroblasts were isolated from each scar tissue approximately two months after injury (Fig. 1a). Because fibroblasts were obtained from injured skin tissues, myofibroblasts were also present in the population, and we identified them with alpha-smooth muscle actin (α-SMA) staining. After two days of culturing the fibroblasts, total RNA was extracted from each group for RNA sequencing (RNA-Seq) (Fig. 1b). The average log_2_(normalized read count (RC)) values for all genes were plotted on the 2D plane (Fig. 1c). The correlation coefficients for total gene expression between different types of fibroblasts were similar (0.97, 0.98, and 0.97 for normal skin vs. hypertrophic scar, hypertrophic scar vs. keloid, and normal skin vs. keloid, respectively), indicating that the total gene expression pattern is incapable of distinguishing fibroblasts from different scar types.

**Fig. 1.**
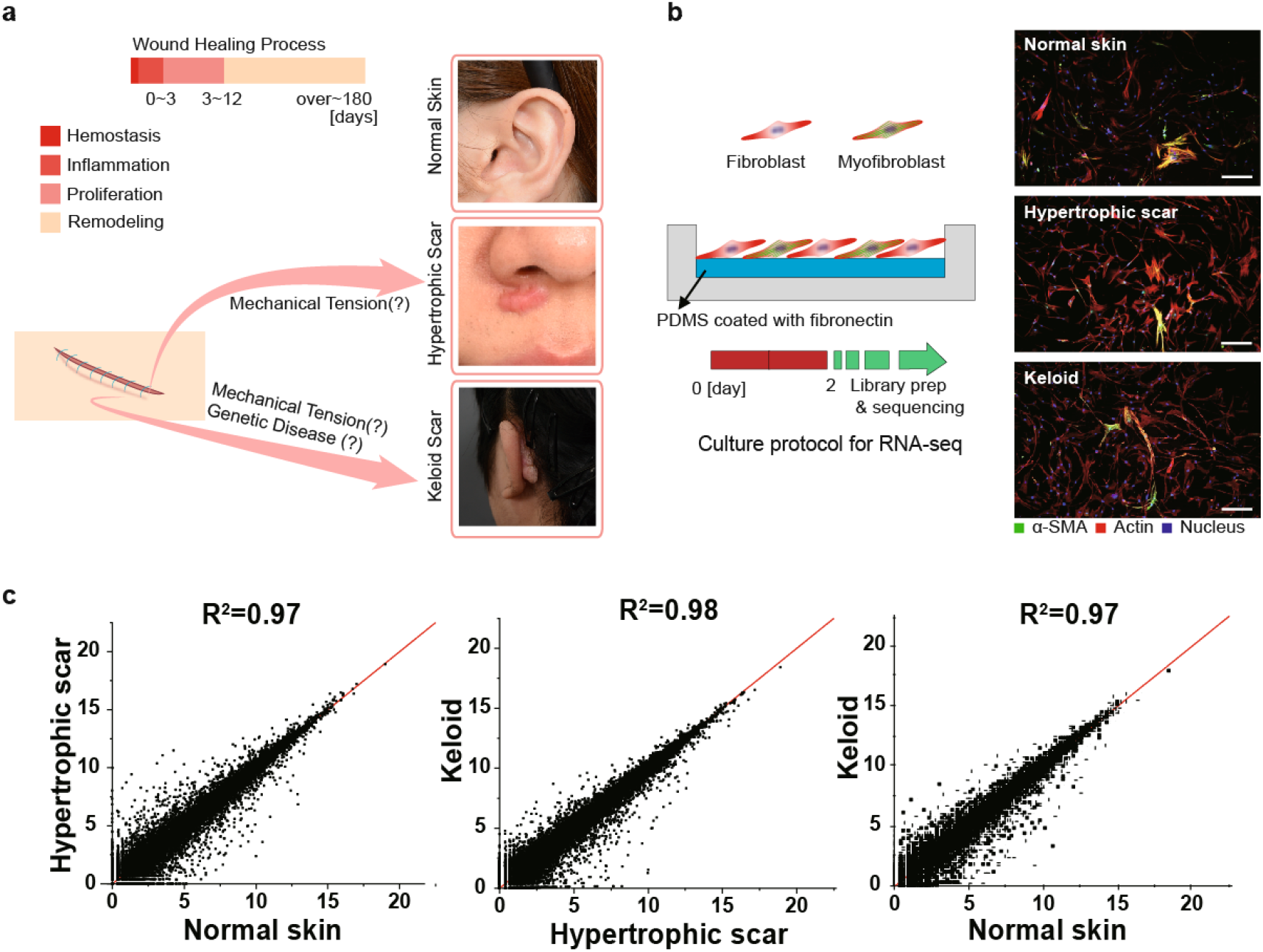
Fibroblasts from normal skin, hypertrophic scars, and keloids and their similar transcriptomic features. (a) Comparison of normal skin, hypertrophic, and keloid scars and their putative causes of formation. (b) Isolation of (myo)fibroblasts from each type of scar tissue and culture protocol for RNA-seq. The scale bar represents 100 μm. (c) The average log_2_(normalized RC) values for total gene expression plotted on the 2D plane and R-squared values for normal skin vs. hypertrophic scar, hypertrophic scar vs. keloid, and normal skin vs. keloid fibroblasts. RC: read count.

Regarding morphology, significantly different features were observed (Fig. 2). In particular, fibroblasts from hypertrophic scars showed significant differences in cell area and aspect ratios compared to those of fibroblasts from normal skin and keloids. Fibroblasts from hypertrophic scars had larger cell sizes and lower aspect ratios. These different morphological features might arise due to the fact that hypertrophic scar tissue contains more myofibroblasts than normal skin and keloid scar tissues. In contrast, fibroblasts from normal skin and keloids showed no significant differences in cell area or aspect ratios.

**Fig. 2.**
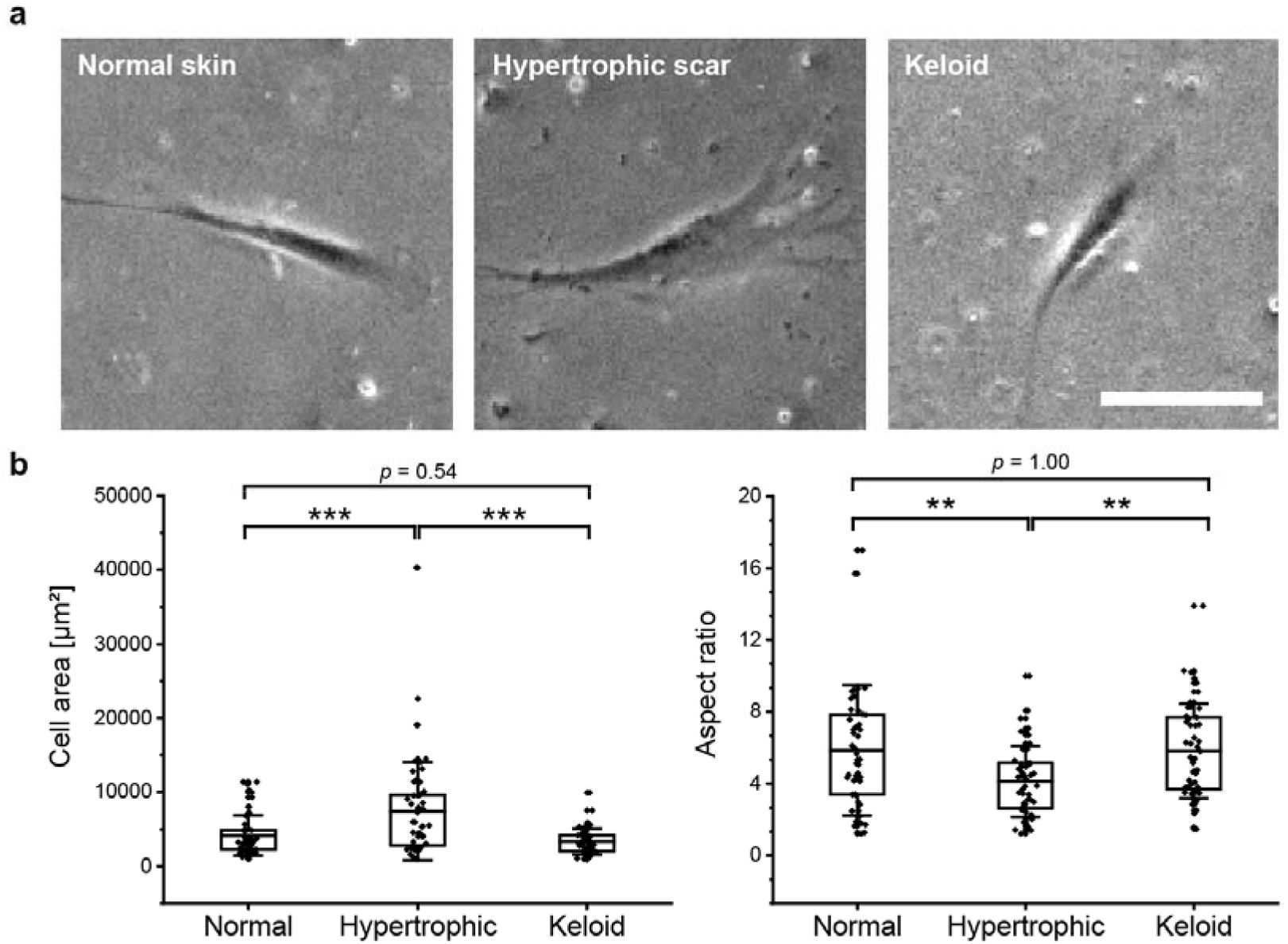
Morphological features of fibroblasts from normal skin, hypertrophic scars, and keloids. (a) Phase contrast microscopy images of fibroblasts from normal skin, hypertrophic scars, and keloids. The scale bar represents 100 μm. (b) Different morphological features of scar fibroblasts. Statistical significance was calculated with one-way ANOVA with Tukey’s post hoc test. \*\**p* < 0.01, \*\*\**p* < 0.001. n=53, 52, and 51 for fibroblasts from normal skin, hypertrophic scars, and keloids. Data represent the mean ± s.e.m.

### Morphogenesis-related *HOX* genes are differentially and specifically expressed in fibroblasts from different scar types

A total of 219 differentially expressed genes (DEGs) were sorted with ExDEGA software with FC (fold change) > 2, log_2_ (normalized RC) > 4, and p < 0.05 as the thresholds (Table S1). For those 219 DEGs, principal component analysis (PCA) was carried out, and the data were projected onto the first two principal components (PC1 and PC2), which accounted for 43.95% and 23.60% of the total variability, respectively (Fig. 3a). Samples from the same scar types clustered together, implying that the DEGs are representative enough to distinguish fibroblasts from each scar type.

**Fig. 3.**
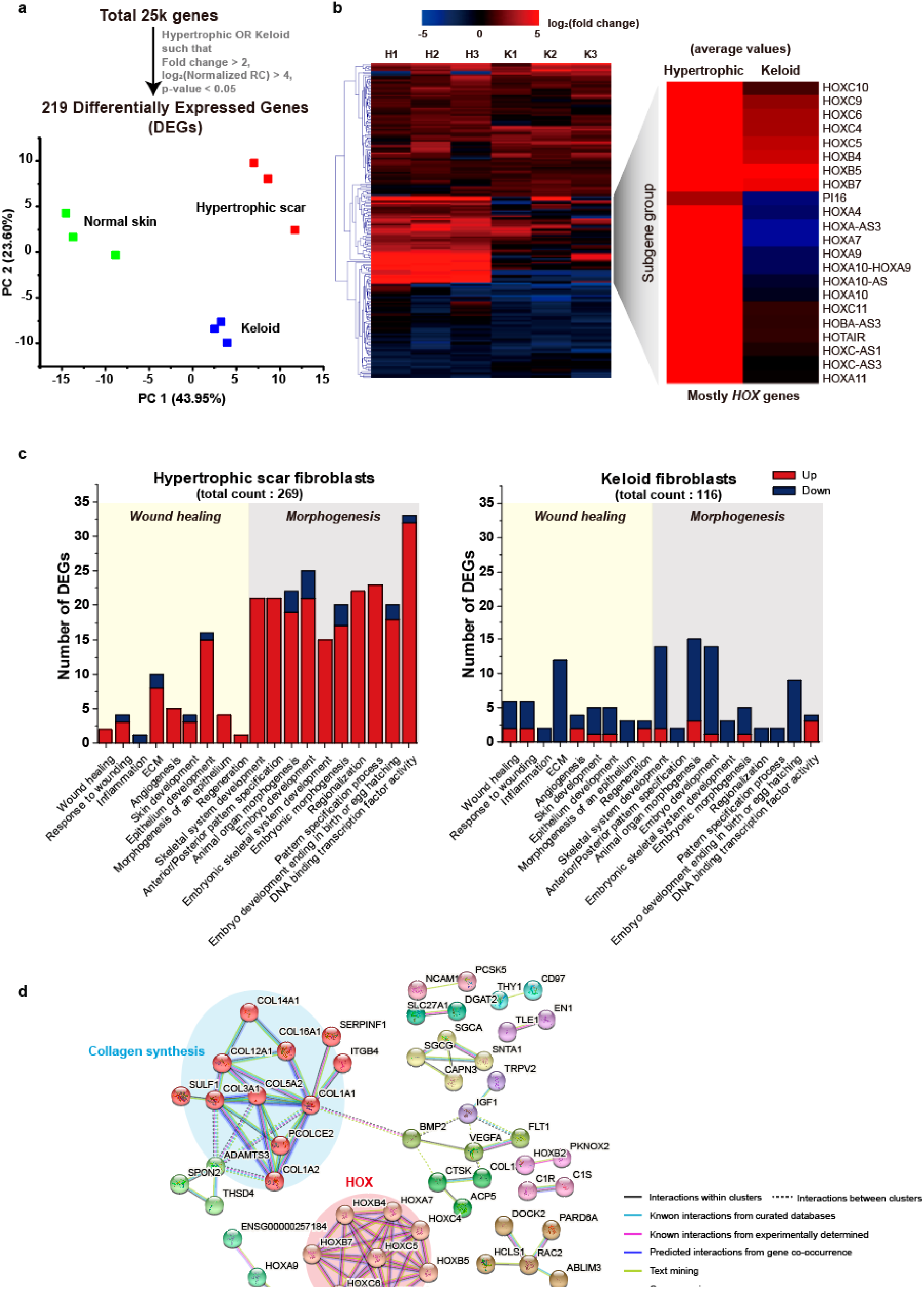
Differentially expressed genes (DEGs) demonstrate the different morphogenetic features of fibroblasts from normal skin, hypertrophic scars, and keloids. (a) Principal component analysis (PCA) of 219 DEGs plotted on the 2D domain (first and second principal components). n = 3 for fibroblasts from normal skin, hypertrophic scars, and keloids. (b) Hierarchical clustering of log_2_(FC) values and the sub-gene group with average values. FC stands for fold change and is calculated with the normalized read count. N, H, and K indicate fibroblasts from normal skin, hypertrophic scars, and keloids; the numbers represent different patient groups. (c) Gene Ontology (GO) analysis of DEGs with wound healing-and morphogenesis-related biological processes. (d) Markov clustering of DEGs using STRING. Network analysis revealed two distinct clusters, collagen synthesis and *HOX* genes. The nodes represent the proteins encoded by the DEGs. Only connected nodes are shown, with disconnected nodes containing 165 genes removed from view.

The hierarchical clustering analysis represented by a log_2_(FC) heatmap revealed a distinct subgene group, which was significantly differentially expressed between fibroblast from hypertrophic scar and fibroblasts from keloid (Fig. 3b). Most of the genes were *HOX* genes, which are responsible for morphogenesis during embryonic development. These genes were highly upregulated in fibroblasts from hypertrophic scar but not in fibroblasts from keloid. In gene ontology (GO) analysis, we selected nine biological processes related to wound healing and the top ten overlapping processes calculated with Molecular Signature Database v7.0, which represented morphogenesis-related biological processes (Fig. 3c). As expected from the heatmap, morphogenesis-related processes were more strongly correlated with fibroblasts from hypertrophic scar than wound healing processes and were mostly upregulated compared to their expression levels in fibroblasts from normal skin. This result may have occurred because the cells were obtained during the last stage of the wound healing process when the wound healing-related genes must have already been downregulated. However, no dominant class was found in fibroblasts from keloid, where most of the genes seemed to be inactivated or downregulated. These data suggest that morphogenesis-related biological processes significantly affect hypertrophic scar formation. On the other hand, keloid formation does not seem to be affected by either of these processes, as evidenced by the fact that the total gene counts for the processes were smaller than those in fibroblasts from hypertrophic scar. (Fig. 3c). STRING analysis with Markov clustering revealed two significant clusters, collagen-encoding genes and *HOX* genes, along with 12 small groups (Fig. 3d). Interestingly, there were no known interactions between the two clusters.

### Wound sites feature injury-induced tension loss in the tissue

Skin tissues are known to be under natural tension during muscle movement and respiration. Therefore, these tissues are in tensional homeostasis. However, tensional homeostasis can be disrupted when external conditions are altered by injury. We first visualized the stress distribution under natural tension and injured conditions by FEM simulation (Fig. 4a). The breach on the top of the stress contour mimics an acute injury to the skin tissue. Excessive tension at the wound site has been thought to cause abnormal scar formation in the form of hypertrophic scars and keloids.

**Fig. 4.**
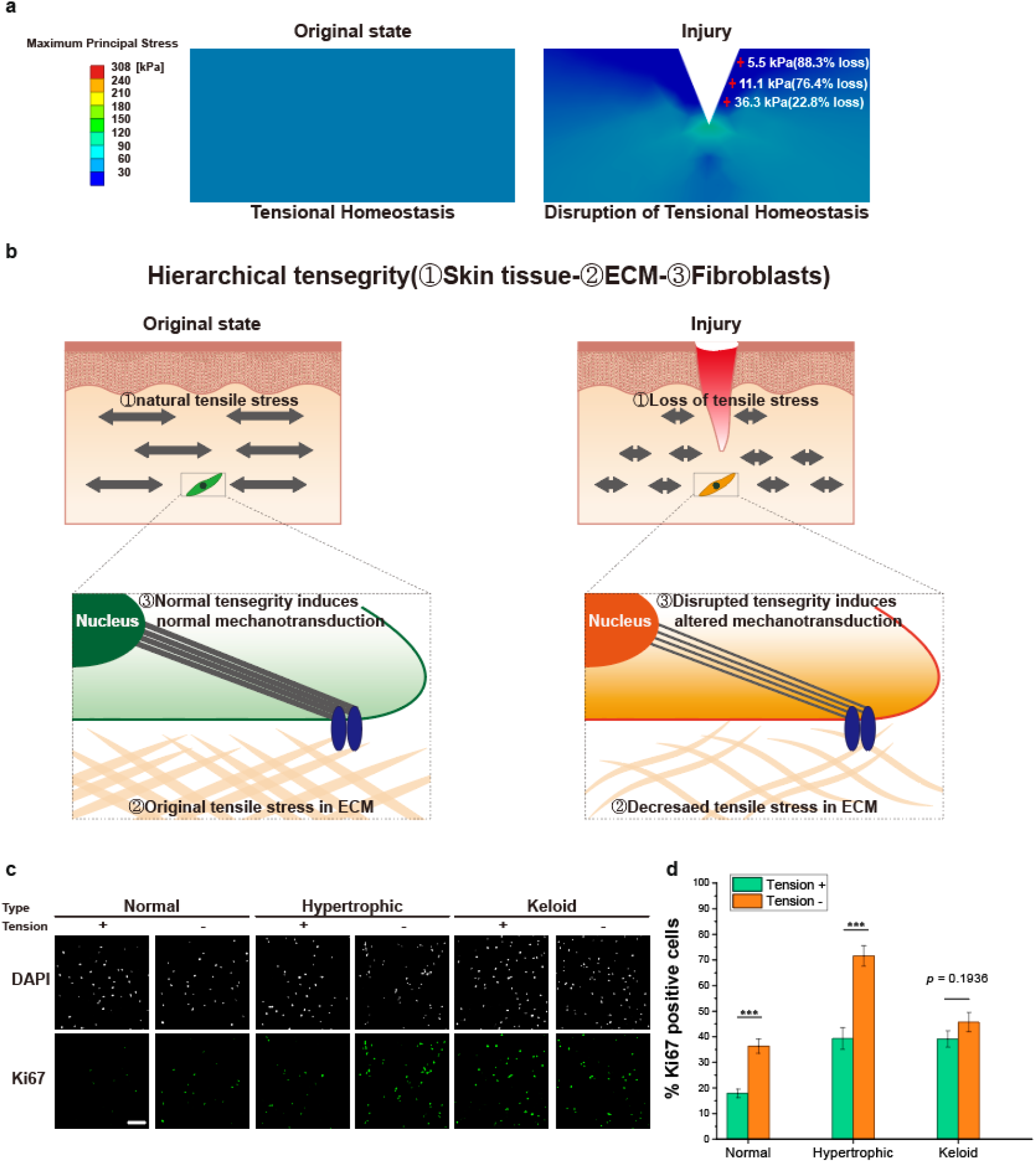
Injury-induced tension loss initiates the wound-healing process by inducing the proliferation of fibroblasts. (a) FEM analysis to visualize the tensile stress distribution in tissues without and with injury. (b) Schematic of hierarchical tensegrity: Skin tissue-ECM-internal cellular tensegrity. (c) Representative Ki67 immunostaining images of fibroblasts from normal skin (left). (d) Proportions of Ki67-positive cells with and without tension. Statistical significance was evaluated with an independent two-tailed t-test. \*\*\**p* < 0.001. The scale bar represents 200 μm. n ≥ 15 images from three independent experiments for fibroblasts from normal skin, hypertrophic scars, and keloids. Data represent the mean ± s.e.m.

In contrast to this common belief, the maximum principal stress was decreased in most areas near the injury. The average principal stress value was 51.1 kPa for the original state and 41.6 kPa for the injury model. Interestingly, the skin showed significant tension loss near the injured area even though there was also a stress concentration effect at the periphery of the injury. We highlighted three locations on the stress distribution map of the injury model with a red cross and investigated the differences in the stress values (Fig. 4a). Compared to those of the identical positions in the original state, the maximum principal stresses were decreased by up to 94.0% in the injured state. This result implies that fibroblasts near the site of injury can experience a dramatic change in their mechanical environment. As mechanical forces influence the wound healing process, the loss of tension upon injury can result in orchestrated responses of fibroblasts^26–28^.

Tensegrity has been proposed to describe how mechanical forces regulate cellular biochemical systems^29–31^. To describe this orchestrated phenomenon, we can introduce the concept of hierarchical tensegrity (Fig. 4b). This concept depicts the hierarchy of tensional integrity transmitted from the skin tissue to the ECM and cells. In other words, cells under natural tension are capable of sensing their mechanical environment and developing stable tensegrity, which results in corresponding biological responses via mechanotransduction. However, when the mechanical environment is compromised by injury, cells can sense the ECM’s loss of tension and adapt their tensegrity, eliciting distinct biological responses.

### Tension correlates negatively with proliferation in fibroblasts from normal skin

We examined the proliferation of fibroblasts in different types of scars, which is a critical feature of the early stage of the wound healing process, as an indicator of altered biological response. To simulate skin tissue’s natural tension and injury with lost tension, we utilized tension and no-tension conditions, respectively, in our tensile stimulation experiment. Immunofluorescence images in Figure 4c revealed Ki67-positive cells among fibroblasts from normal skin, indicating that the fibroblasts without tension had a higher number of Ki67-positive cells than those under tension, which suggests that tension suppresses proliferation of fibroblasts from normal skin. However, looking at the experimental results from a reverse logic perspective, we found a negative correlation between tension and proliferation, suggesting that the loss of tension in the tissue following an injury could result in increased proliferation. Therefore, we may consider that tension loss induced by injury could serve as a contributing factor to stimulate fibroblasts for wound healing, leading to an increase in their proliferation.

We also quantified the percentage of Ki67-positive cells among fibroblasts from normal skin and fibroblasts from other types of scars (Fig. 4d). The graph shows that fibroblasts from normal skin and hypertrophic scar showed a significantly higher percentage of Ki67-positive cells in the absence of tension conditions. However, regardless of the tension conditions, the population of fibroblasts from keloids exhibited no differences in proliferation and maintained a high level of proliferation. Our experimental findings corroborated that fibroblasts from keloids feature hyper-proliferation ^32^.

### Tension positively correlates with the expression of *HOX* genes and collagen-encoding genes in fibroblasts from normal skin

We discovered that *HOX* genes, which are critical players in morphogenesis, were differentially expressed between scar types using RNA-Seq. We performed real-time qPCR with selected *HOX* genes after tensile stimulation to investigate the influence of tension on these genes (Fig. 5a). We selected *HOXA9* and *HOXC10* from the sub-gene group identified through RNA-Seq analysis. *HOXA9* is known to be associated with the wound healing process and hypertrophic scar formation, as reported in previous studies ^33,34^. Meanwhile, *HOXC10* exhibited the highest expression levels in fibroblasts from hypertrophic scars compared to those from normal skin and keloid. Two-way ANOVA revealed that only one main factor, tension, significantly affected the expression levels of *HOXA9* and *HOXC10* (Table S2, 3). In fibroblasts from normal skin, these two *HOX* genes were downregulated in the absence of tension compared to the tension group. Hypertrophic scar fibroblasts also showed a similar tendency, but the expression levels were still higher than normal levels under both conditions. Interestingly, *HOX* gene expression levels in fibroblasts from keloid did not statistically differ between the tension and non-tension conditions, which is consistent with the proliferation trend of this population.

**Fig. 5.**
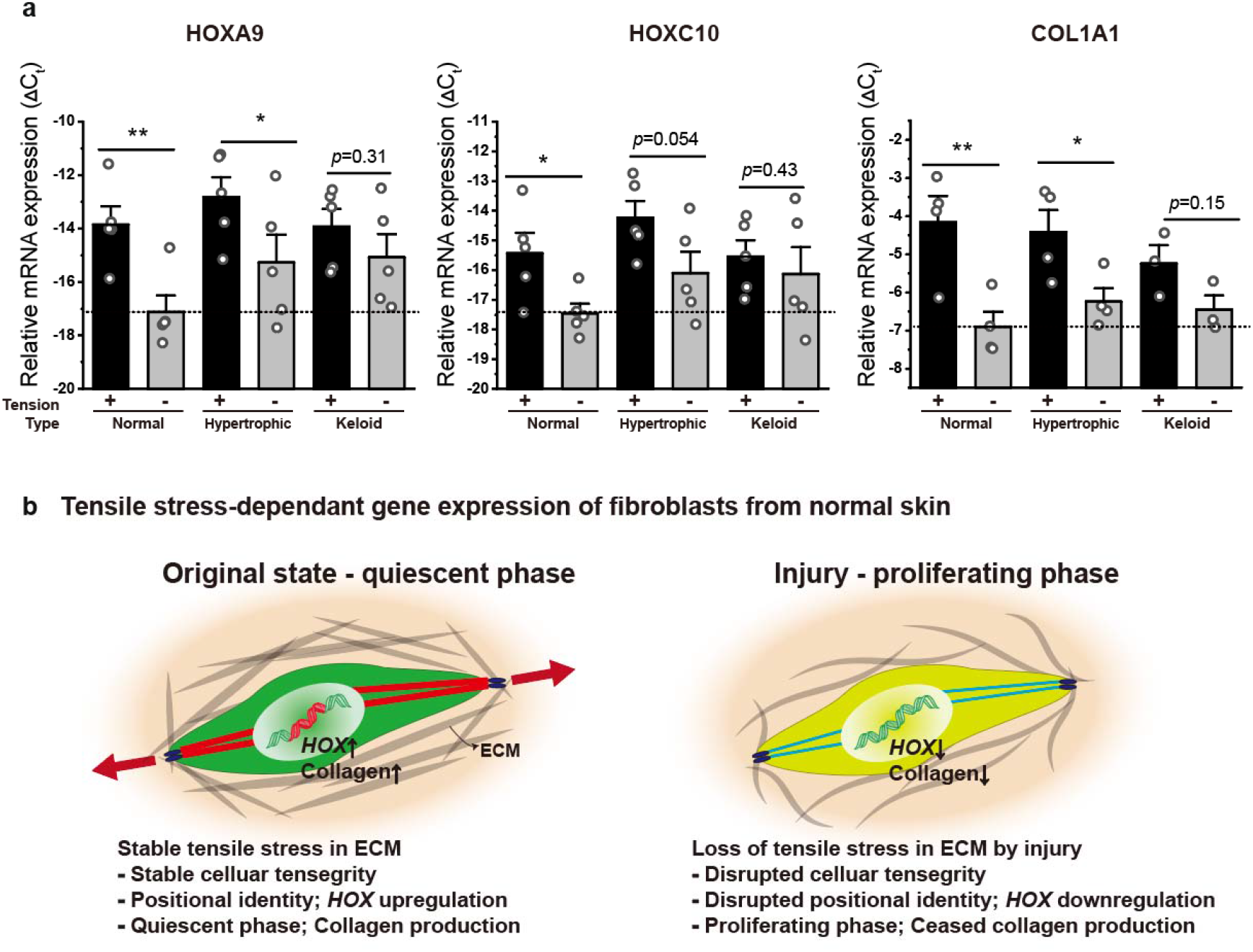
Positive correlation between tension and selected genes (*HOXA9*, *HOXC10*, and *COL1A1*). (a) *HOXA9, HOXC10,* and *COL1A1* relative mRNA expressions of fibroblasts from normal skin, hypertrophic scars, and keloids. + and – represent with and without tension, respectively. Statistical significance was evaluated via two-way ANOVA with post hoc tests (*HOXA9* and *HOXC10*: n=5. *COL1A1*: n = 4, 4, and 3 for fibroblasts from normal skin, hypertrophic, and keloids). \**p* < 0.05, \*\**p* < 0.01. Data represent the mean ± s.e.m. (b) Schematic of the genetic response to tension in fibroblasts from normal skin. Fibroblasts from normal skin express *HOX* and collagen synthesis genes in their normal state. However, when the existing tension is lost due to injury, the expression of these genes is decreased via mechanotransduction.

Additionally, we performed qPCR on the collagen synthesis gene *COL1A1*. Rognoni *et al*. reported that collagen synthesis and fibroblast proliferation were inversely related^35^. The *COL1A1* gene expression level was decreased in fibroblasts from normal skin and hypertrophic scars in the absence of tension, a condition that promotes proliferation. However, fibroblasts from keloid scars did not exhibit a mechanosensitive response to tension regarding *COL1A1* expression (Table S4).

Assuming that the reverse logic holds true, implying that a loss of tension enhances the proliferation of fibroblasts from normal skin, we can integrate proliferation with gene expression patterns to suggest a hierarchical tensegrity-based response to tension (Fig. 5b). The left image depicts the original state of fibroblasts from normal skin and their biological responses to tension. They establish and maintain stable cellular tensegrity under natural tension. Therefore, *HOX* genes are expressed in adult cells as positional identity markers^36,37^. Because the cells are in a quiescent state, their collagen synthesis gene expression is upregulated. However, if an injury compromises the skin’s integrity, fibroblasts can sense the loss of tension and alter their response via hierarchical tensegrity. As a result of the disruption of positional identity, *HOX* genes are downregulated, and collagen synthesis is temporally ceased, which allows the cells to proliferate. These findings suggest a plausible link between *HOX* and collagen synthesis-related gene expression, but no direct evidence has been shown thus far (Fig. 3d).

## Discussion

The etiology of hypertrophic scars and keloids remains unclear. While many investigations have suggested that keloids have a significant hereditary susceptibility, especially in those with dark skin, it is essential to recognize that this genetic predisposition may still lack a clear causal link. This study showed distinct expression levels of *HOX* genes in fibroblasts from different scar types, whose mechanosensitivity to exogenous tension varied markedly.

Previous studies have reported differential expression of *HOX* genes in cells from normal skin and keloids^38,39^. Specifically, cells from keloids showed lower expression of multiple *HOX* genes such as *HOXA7*, *HOXA9*, *HOXC8*, and *HOXC10* compared to normal fibroblasts. However, Xie et al. reported that the overexpression of *HOXB9* could facilitate hypertrophic scar formation^40^. These studies have focused on the expression of specific *HOX* genes in the hypertrophic scar or keloid formation. In contrast, our study compared the whole gene expression of cells from normal skin, hypertrophic scars, and keloids rather than focusing on specific *HOX* genes for analysis. We then identified 219 DEGs related to morphogenesis-related biological processes, which were representative enough to distinguish between each sample type (Fig. 3a). Among those 219 DEGs, we identified a specific subgene group consisting mostly of highly upregulated *HOX* genes in cells from hypertrophic scars but not in cells from keloids in our data (Fig. 3b). Our data-driven analysis suggests that the subgene group found is consistent with previous findings about differential *HOX* gene expression in fibroblasts from different scar types.

Physical factors like excessive tension have been thought to influence hypertrophic scar and keloid formation. This is because most of those scars are found in regions with relatively high tension (the neck, chest, and similar sites), and patients can also feel the unpleasant tension around the wound. To our surprise, this study demonstrated that the tension near the wound site decreased dramatically based on FEM analysis.

From the viewpoint of solid mechanics, skin, a material in natural tensional homeostasis, experiences a decrease in tensile stress upon injury. In addition, because of the skin’s residual stresses, the injury’s periphery can be pulled outward, making the wound larger. This phenomenon can be viewed as tension acting on the wound, even though there is no active tension on the material (i.e., skin).

Based on our results, we suggest a significant contribution of hierarchical tensegrity on scar formation. In short, tension can be transmitted from the tissue level to the ECM and finally to the constituent cells. Fibroblasts establish appropriate cellular tensegrity when the tension is transmitted, leading to differential mechanotransduction. Therefore, if fibroblasts sense a loss of tension, they would exhibit altered cellular tensegrity, inducing different biological processes. The altered biological response to the loss of tension can be a critical factor in wound healing, and scar formation and clinical observations support this idea. Consistent observations made by dermatologists and plastic surgeons indicate that wounds of elderly individuals tend to heal with thinner scars in comparison to younger patients ^41,42^. While the rate of collagen production decreases with aging and the healing in the elderly was thought to be defective, there is an agreement that healing in the aged is delayed, but the ultimate result is qualitatively identical to that in young subjects^43^. With this perspective, we can see that the low-tension environment induced by the compromised architecture of old skin may help fibroblasts from older individuals to establish their normal cellular tensegrity. This could explain why fibroblasts in older individuals would not experience a dramatic decrease in tensile stress when injured, resulting in reduced scar formation. Our idea was corroborated by another report in which fetal mouse skin tissue was shown to have low resting tension and not likely to develop significant scars^44^.

Previous studies have demonstrated the influence of mechanical stimulation on differentiation and proliferation in vitro^45–47^. These findings suggest that a specific range of mechanical tension may function as a morphogenetic cue for local tissue pattern formation in vivo. In terms of fibroblast-to-myofibroblast differentiation, it has been established that mechanical stimuli play a crucial role in increasing ECM proteins and proteoglycan content^6,26,46,48^. In addition, the effect of cyclic mechanical stimulation on fibroblast proliferation has been extensively investigated, revealing its ability to either enhance or inhibit proliferation^49–53^.

While we acknowledge the effect of mechanical cues on fibroblast differentiation, our focus was on the impact of tension on fibroblast proliferation, as excessive proliferation is one of the primary features of pathological scar formation^54,55^. We observed a negative correlation between tension and proliferation in tensile stimulation experiments. By applying reverse logic to this finding, we were able to establish the hypothesis that if there is tension loss in the skin, fibroblasts will proliferate more actively, which implies initiation of the proliferative phase of the wound healing process. Based on this hypothesis, we set an external tension condition as a control condition to mimic natural tension in humans and a no-tension condition as an injury-mimicking condition for the in vitro tensile stimulation experiment. The selected *HOX* genes were downregulated in the absence of tension in fibroblasts from normal skin and hypertrophic scars, even though the expression levels of hypertrophic scar fibroblasts were still higher than those observed under normal conditions. However, fibroblasts from keloids had similar or lower expression levels compared to fibroblasts from normal skin and hypertrophic scars under tension conditions and showed no statistically different expression pattern changes in the absence of tension. *COL1A1* gene expression patterns also showed a similar tendency. *COL1A1* gene expression patterns also showed a similar tendency. This implies that collagen synthesis and *HOX* expression are positively correlated in fibroblasts from normal skin and that a pathway exists between them, even though no such interactions have been reported thus far (Fig. 3d). From these experimental results 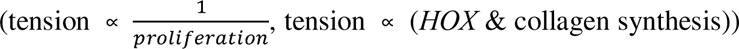, one can predict that fibroblasts under natural tension conditions will proliferate less and upregulate collagen synthesis gene expression levels compared to fibroblasts under the injury-induced loss of tension condition. However, contradictory responses of fibroblasts to tension have been reported by other researchers^50,52,53,56–59^. We believe these discrepancies originate from differences in stimulation conditions, including differences in strain, frequency, duration of stimulation, and resting period, as well as other conditions of the experimental setup, such as the rigidity of substrates, ECM type, and ECM concentration, resulting in differential activation of mechano-responses in fibroblasts^60^.

Overall, we provide a new viewpoint on wound healing and scar formation scenarios in which the successful wound healing process requires homeostatic maintenance of natural tensile stress, as illustrated in Fig. 6a (adapted from Rognoni *et al*.^35^). This tensional homeostasis is strongly correlated with *HOX* gene expression in fibroblasts. Injury threatens tensional homeostasis because the loss of tensile stress is transmitted to resident cells through hierarchical tensegrity. Upon the loss of tension, quiescent fibroblasts become activated and start proliferating. When the fibroblast population reaches a sufficient size, the fibroblasts exit the cell cycle and return to a quiescent state in which cells efficiently deposit ECM for remodeling^35^. Based on the negative feedback loop of proliferation-ECM deposition suggested by Rognoni *et al*., we can speculate that excessive deposition of ECM can lead to scar formation.

**Fig. 6.**
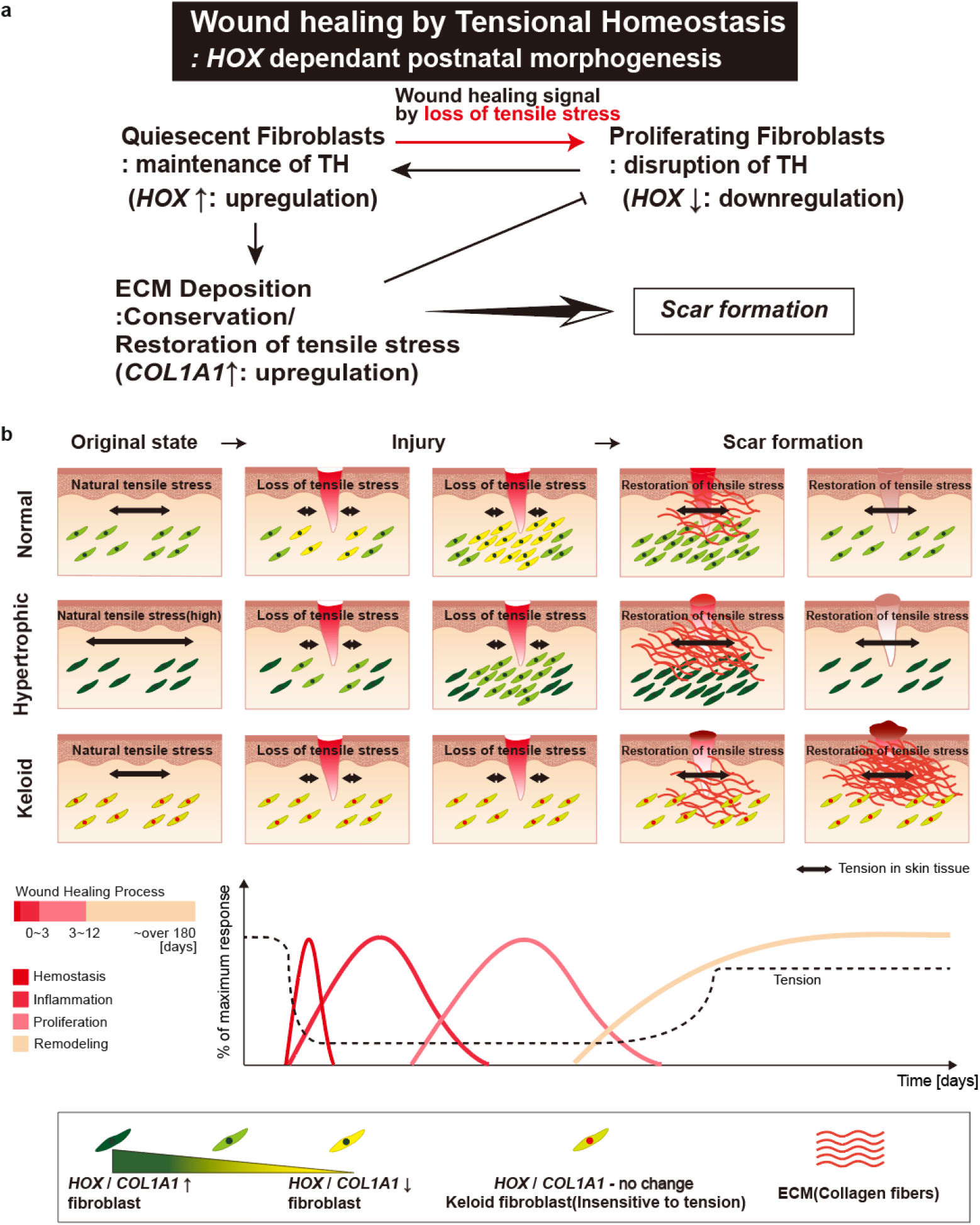
Novel wound healing model and scar formation scenarios that include mechanical tension and genetic effects. (a) Schematic of the tensional homeostasis-driven wound healing process. The schematic was adapted from Rognoni *et al*.^35^, CC, by 4.0 (https://creativecommons.org/licenses/by/4.0/). TH: tensional homeostasis. (b) Schematic diagram of normal, hypertrophic, and keloid scar formation scenarios with respect to tension-sensitive *HOX* gene expression.

These processes can be orchestrated by the expression of *HOX* genes, which encode critical transcription factors for morphogenesis and the positional memory of fibroblasts. It has been suggested that fibroblasts have a program to maintain their positional memory^37^. Therefore, decreased expression of *HOX* can be viewed as a threat to this maintenance program, resulting in systematic responses to restore the program. We view this process as *HOX*-dependent postnatal morphogenesis, in which fibroblasts restore the original dermal architecture through the regulation of *HOX* expression levels.

This wound-healing model aligns well with conventional scar treatments, which have been used based on empirical evidence. For example, silicon patches and force-modulating tissue bridges are used to prevent severe scar formation^61^. Pressure therapy for hypertrophic burn scars is also effective in both the prevention and treatment of scar formation^62^. All these treatments can be viewed as strategies to recover the original tensional state of the wound site.

With the suggested model, we were able to differentiate hypertrophic scars from keloids. Any abnormalities during the orchestrated healing process can lead to scar formation, as depicted in the plausible scenarios shown in Fig. 6b. For example, hypertrophic scars can form when the cells in high-tension regions maintain a high level of *HOX* gene expression upon wounding, maintaining an active ECM deposition phase during the wound healing process. On the other hand, keloids can be formed by fibroblasts, which are consistently proliferating regardless of tension and seem to be inherently insensitive to the loss of tension. Therefore, other cells regulating the ECM balance will be damaged when an injury compromises the skin tissue. Then, these fibroblasts continue depositing ECM externally, resulting in keloids that push against the original wound boundaries.

It is plausible that fibroblasts from keloid exhibit an intrinsic dysfunction of *HOX* gene expression in response to injury-induced tension loss, and this intrinsic issue related to *HOX* gene expression might be regarded as genetic susceptibility in the field. To our knowledge, the suggested wound healing model and scar formation scenarios are the first demonstration of the mechanisms by which different hypertrophic and keloid scars can form in terms of mechanical cues and gene expression.

However, there are several limitations to our study. Firstly, it was impossible to acquire fibroblasts from the same age group, sex, and body parts. Therefore, further investigations of *HOX* genes and similar patient conditions (in terms of age, sex, and injury site) will help strengthen our model and improve the understanding of scar formation. Also, because in vitro experiments cannot fully recapitulate the long-lasting and complex wound healing phenomena, our in vitro tensile stimulation experiments were designed to maximize the effect of tension on the cells. Therefore, even though we believe that the time frame set up in our experimental design was appropriate to observe the different behaviors of cells upon tensile stimulation, one should be noted that there could be remodeling and changes to mechanics that occur at the wound site in vivo, which could influence the wound healing process. In addition, while *HOX* gene expression patterns (*HOX* code) are known to be site-specific, which allows prediction of the cells’ origins^63,64^, our tensile stimulation experiments only evaluated two *HOX* genes, *HOXA9* and *HOXC10*. Although the fundamental mechanism of tension-induced *HOX* gene regulation requires further investigation, our findings elucidate the etiology of hypertrophic scar and keloid formation. A better understanding of *HOX* gene expression and scar formation will open the door to new therapeutic strategies for scarless wound healing.

## Methods

### Primary cell isolation and culture

Three normal skin tissues, three hypertrophic scar tissues, and three keloid scar tissues were obtained from plastic and reconstructive surgery patients, and the information about donors were described in Table S5. Hypertrophic scars and keloids were diagnosed by plastic surgeons. Briefly, scars that have not overgrown the original wound boundaries are defined as hypertrophic scars, scars that have overgrown the original wound edges are defined as keloids, and some samples were histologically verified. Before surgery, all patients were informed of the purpose and procedure of this study, and the patients agreed to donate excess tissue. In addition, written informed consent was obtained from all participants or their legal representatives. This study was approved by the institutional review board (IRB) of KAIST (KH2017-75) and performed.

Patients’ normal skin, hypertrophic scar, and keloid scar tissue were washed several times with phosphate-buffered saline (PBS, pH 7.4) and incubated with 2.4 units of Dispase® (Sigma, St. Louis, MO, USA) for 12 h. After washing with PBS, the dermis and epidermis were separated using forceps. Next, the separated dermal tissue was cut into small pieces and incubated with 0.2% collagenase type II (Worthington Biochemical Corp. Lakewood, NJ. USA) for 1 h, and the cells were precipitated using centrifugation at 1000 rpm for 5 min. After several washes, the cell suspension was filtered through a cell strainer (pore size 75 μm, Nunc, Roskilde, Denmark). The filtered cells were grown in Dulbecco’s modified Eagle’s medium (DMEM, Gibco/BRL, Gaithersburg, MD, USA) containing 100 units/mL penicillin G, 100 μg/mL streptomycin sulfate (Sigma), and 10% fetal bovine serum (Gibco/BRL) in 100-mm culture dishes in an incubator at 37 L and 5% CO_2_. Once the cells had reached 70– 80% confluence, they were treated with Trypsin 0.25% (1x) solution (Hyclone Laboratories Inc., Logan, UT, USA) for passage.

### Antibodies and Immunofluorescence microscopy

Cells were washed with PBS and fixed for 15 min in 4% (v/v) paraformaldehyde. After washing three times with PBS, cells were permeabilized for 20 min in 0.2% or 1% (v/v) Triton-X (Sigma Aldrich) and blocked for 60 min in 3% (v/v) bovine serum albumin (BSA). After washing in PBS, cells were incubated with diluted primary antibodies (α-SMA and Ki67, 1:100, Abcam) for 12 hours and secondary antibodies (Alex Flour 488 and 568-fluorescence, 1:200, Invitrogen) for 1 hour at room temperature. To label nuclei, 4’,6-diamidino-2-phenylindole (DAPI, 1:50000 dilution, Molecular Probes) was treated in the cells for 3min. Cells were then imaged using multichannel fluorescence microscopy (Carl Zeiss).

### NGS and data analysis

Cells were cultured on polydimethylsiloxane (PDMS) (Dow Corning, USA), which were pretreated with oxygen plasma (Convance, Femto Science, Korea) for 1 min at a pressure of 0.7 Torr followed by 1μg/cm2 of fibronectin (Invitrogen, USA) coating for 120 min at room temperature. Once the cells had reached 90% confluence, cells were lysed in situ, and total RNA was extracted using a Trizol reagent (Invitrogen, USA). RNA purity was assessed by Agilent 2100 bioanalyzer with the RNA 6000 Nano Chip (Agilent Technologies, Amstelveen, The Netherlands), and RNA was quantified at 260 nm with ND-2000 Spectrophotometer (Thermo Inc., DE, USA).

We generated libraries using QuantSeq 3’ mRNA-SeqLibrary Prep Kit (Lexogen, Inc., Austria) according to the manufacturer’s instructions. Briefly, each 500ng total RNAs were reverse transcribed using an oligo-dT primer containing an Illumina-compatible sequence at its 5’ end. After the degradation of the RNA template, the second-strand synthesis was commenced with a random primer. The random primer contains an Illumina-compatible linker sequence at its 5’ end. Magnetic beads were used for the purification of the double-stranded library. Then, amplification of the library was carried out for cluster generation. The obtained library is purified from the PCR products and sequenced using NextSeq 500 (Illumina, Inc., USA) to produce 75 base pair single-end reads. QuantSeq 3’ mRNA-Seq reads were mapped to the human genome using Bowtie2 (Langmead and Salzberg, 2012). The alignment file was used to assemble transcripts, estimate their abundances and detect differential expression of genes. The quantile normalization method was used to process the RC (Read Count) data using EdgeR within R (R Development Core Team, 2016) using Bioconductor^65^. Differentially expressed genes (DEGs) were filtered with ExDEGA software (ebiogen, Korea). Hierarchical clustering was performed with MultiExperiment Viewer(MeV, version4.9.0)^66^. Euclidean distance was set for the distance metric, and average linkage clustering was set for the linkage method. Biological process classification was based on searches done by Gene Ontology annotation and Molecular Signatures Database v7.0 by Broad Institute. Network analysis was conducted using STRING version 11.5^67^. The STRING network analysis parameters were set to all the sources with a high confidence score (0.7). Markov clustering with an inflation parameter setting of 3 was used to visualize the network analysis.

### Finite element method (FEM) analysis for wounded skin tissue

FEM analysis was conducted to see the difference in stress distribution with the commercial code ANSYS-Mechanical. The mechanical properties of skin are very heterogeneous not only among the different layers but also with anatomical skin region as well as other factors such as age, sex, and pathological condition^68^. Though the correct value of Young’s modulus is a critical property for FEM simulation, the values may differ depending on skin locations or the test methods used for the measurement. We set Young’s modulus of 1 MPa for our simulation, which was measured by the uniaxial tensile test on the forearm and is similar to Young’s modulus of the PDMS chamber that we used for our experiments ^69,70^. Moreover, the Poisson ratio was set to 0.48^71^. It is known that under physiological conditions, fibroblasts in skin tissues experience 4-10 % strain^72^. Therefore, we imposed a 5 % strain, which is the same percent strain used in our experiments, to the one wall and set fixed support conditions for the confronting wall for boundary conditions to simplify the problem.

### Application of a mechanical tensile stimulus to fibroblasts

For the tensile stimulation experiments, fibroblasts from the last donor group were used (third donor of normal skin, hypertrophic scars, and keloid, respectively, Table S5). Fibroblasts were cultured on PDMS chambers we designed (the same PDMS used in NGS analysis), which were pretreated with oxygen plasma for 1 min at a pressure of 0.7 Torr followed by 1μg/cm^2^ of fibronectin coating for 120 min at room temperature. The PDMS chambers with fibroblasts were loaded on a lab-made uniaxial cyclic tensile stimulation device (Fig. S1). Because the fibroblasts were attached to the fibronectin-coated-PDMS chambers and cells produce other matrix proteins during the pre-incubation period, the attached cells can be stretched by applying tension on a flexible membrane via a variety of ligand/receptor interactions, including integrin linkages at focal adhesions. For physiological relevance, intermittent stimulations were used rather than exhaustive continuous stimulation, where we applied six cycles of intermittent tensile stimulation to the fibroblasts (5% strain, 0.5Hz), where each cycle consisted of a 10-minute stretching followed by a 60-minute resting period. All stretch experiments were carried out inside an incubator at 37°C in 5% CO_2_. Unstretched cells were incubated under the same conditions as the samples undergoing cyclic tension.

### Reverse transcription Quantitative PCR

At the end of tensile stimulation experiments, total RNA was isolated from fibroblasts with RNAiso reagent (Takara Bio, Japan) according to the manufacturer’s instructions. Extracted RNAs were reverse transcribed to cDNA using iScript cDNA Synthesis Kits (Bio-Rad, USA) and Biometra T-personal Thermal Cycler for the synthesis. Real-time qPCR was performed in duplicates with iQ SYBR green supermix (Bio-Rad, USA) and a Bio-Rad CFX96 real-time detection system. Glyceraldehyde 3-phosphate dehydrogenase (GAPDH) was used for the reference gene. ΔC_t_ values were used for hypothesis testing and used to express relative mRNA expression. ΔC_t_ = C_t_(reference gene) - C_t_(gene of interest), C_t_: Threshold cycle

The following primers were used:

GAPDH (For: CTGGGCTACACTGAGCACC, Rev: AAGTGGTCGTTGAGGGCAATG), HOXA9(For: CTGTCCCACGCTTGACACTC, Rev: CTCCGCCGCTCTCATTCTC), HOXC10 (For: CTATCCGTCCTACCTCTCGCA, Rev: CCTGCCAACAGGTTGTTCC) COL1A1(For: GTGCGATGACGTGATCTGTGA, Rev: CGGTGGTTTCTTGGTCGGT).

### Statistical analysis

Data are expressed as mean ± s.e.m., unless otherwise stated, and the number of biological and technical replicates is indicated in the figure caption. A two-tailed t-test and ANOVA with post hoc tests was used for hypothesis testing. Differences were considered significant if \**p* < 0.05, \*\**p* < 0.01 and, *** *p* <0.001 after any adjustment. Statistical tests were performed using *jamovi* version 1.6.23.0 (The jamovi project (2021), https://www.jamovi.org).

## Data availability statement

RNA-seq data have been deposited in the Gene Expression Omnibus under accession code GSE210434.

## Acknowledgment

This research was supported by National Research Foundation granted by the Korean Government (NRF-2017R1A2B2007673, NRF-2021R1A2C3008408).

## Author contributions

U.H.K, H.Y.C, and J.H.S conceived the study. M.K., U.H.K. and J.H.S. conceptualized and designed experiments. H.M.K., E.J.O., and H.Y.C. provided the patients’ cells. M.K. and U.H.K. performed the experiments. M.K. and J.H.S. interpreted data and wrote the manuscript. All authors edited the manuscript.

