## SUPPLEMENTARY MATERIALS for "Differential expression of tension-sensitive *HOX* genes in fibroblasts is associated with different scar types"

**Table S1. 219 differentially expressed genes (DEGs) list and expressions.** DEGs were sorted at a FC (fold change) > 2, log_2_(normalized RC) > 4, and p < 0.05 as the threshold. log_2_(normalized RC) values were listed for each N, H, K samples. RC : read count. N, H, and K indicate normal, hypertrophic, and keloid scar fibroblast samples and numbers represent different patient groups.

|  | Gene | N1 | N2 | N3 | H1 | H2 | H3 | K1 | K2 | K3 |
| --- | --- | --- | --- | --- | --- | --- | --- | --- | --- | --- |
| 1 | PHKA1 | 5.51059 | 5.02938 | 5.26706 | 4.84322 | 4.62218 | 4.78806 | 6.45278 | 6.88049 | 6.09714 |
| 2 | ADORA2B | 6.17965 | 6.07176 | 4.88863 | 3.6366 | 4.0374 | 4.35501 | 5.06752 | 5.67881 | 5.92412 |
| 3 | TJP2 | 7.36463 | 8.1942 | 7.8457 | 6.55054 | 6.67388 | 6.07786 | 7.7075 | 7.75051 | 7.96993 |
| 4 | TBC1D31 | 6.47698 | 5.60106 | 5.93597 | 5.02392 | 4.03682 | 4.8405 | 6.34174 | 5.62807 | 5.70071 |
| 5 | SPC24 | 3.04819 | 2.49006 | 2.79 | 3.11019 | 2.19597 | 3.49018 | 3.8054 | 4.59774 | 4.11968 |
| 6 | HOXB4 | 4.13943 | 1.90198 | 0 | 8.68793 | 8.24686 | 7.84056 | 1.1068 | 7.89244 | 0.2842 |
| 7 | C17orf96 | 6.65095 | 6.50052 | 5.88854 | 4.39536 | 5.35925 | 5.35512 | 6.17998 | 6.2933 | 6.29823 |
| 8 | TMEM144 | 4.63361 | 4.42829 | 4.06983 | 3.25999 | 2.87042 | 3.61699 | 4.43052 | 4.23779 | 4.49652 |
| 9 | PRICKLE2-AS3 | 4.95695 | 5.32927 | 5.1114 | 5.18609 | 4.57058 | 5.03225 | 3.57285 | 4.47318 | 3.34328 |
| 10 | PCSK5 | 5.3364 | 6.11089 | 6.11116 | 5.76998 | 5.54122 | 5.7332 | 5.1137 | 4.67595 | 3.66956 |
| 11 | HCLS1 | 2.37315 | 3.48407 | 3.44088 | 7.15577 | 8.04348 | 6.77455 | 7.4084 | 3.35173 | 4.59721 |
| 12 | LOC400043 | 4.22189 | 0.90727 | 4.30334 | 7.46778 | 8.26889 | 7.73353 | 4.36356 | 4.52134 | 7.3571 |
| 13 | ZNF536 | 1.63413 | 1.48943 | 0 | 3.74641 | 5.73612 | 5.20062 | 3.28015 | 4.23963 | 0.23481 |
| 14 | CAPN3 | 5.09507 | 5.48675 | 5.50551 | 5.49209 | 5.59797 | 5.39012 | 4.50581 | 4.5252 | 3.93001 |
| 15 | BAALC | 7.44426 | 4.59949 | 2.56676 | 9.41969 | 10.30028 | 8.99632 | 9.04905 | 6.5865 | 6.65403 |
| 16 | GRP | 3.22158 | 3.9865 | 0 | 6.00185 | 6.85533 | 7.02206 | 4.36361 | 9.87189 | 7.86344 |
| 17 | PLEKHG4B | 2.85838 | 2.2229 | 0.98296 | 3.52204 | 6.14055 | 6.20141 | 4.35957 | 4.23262 | 3.34173 |
| 18 | VAT1L | 2.85828 | 0 | 0.98298 | 4.33026 | 5.89622 | 5.11934 | 4.9176 | 5.68559 | 0.25139 |
| 19 | SPON2 | 9.89748 | 6.86474 | 6.37374 | 12.41042 | 11.30135 | 11.62736 | 12.61374 | 11.05408 | 8.60303 |
| 20 | FLT1 | 4.95889 | 4.06897 | 6.02582 | 8.80649 | 6.67384 | 9.06943 | 7.21328 | 6.87548 | 7.99912 |
| 21 | THRB | 5.47748 | 5.32652 | 4.56652 | 6.4289 | 6.42356 | 6.47571 | 5.57327 | 3.46746 | 5.83523 |
| 22 | HOXD8 | 6.39178 | 3.06979 | 6.17124 | 8.41577 | 8.34658 | 9.29774 | 7.67751 | 6.20942 | 8.18765 |
| 23 | F10 | 3.63634 | 2.70726 | 0 | 5.29501 | 5.62517 | 6.00957 | 3.43636 | 6.23412 | 5.69489 |
| 24 | SYPL2 | 3.63615 | 0.90727 | 2.78898 | 6.10836 | 5.62596 | 5.35426 | 6.04126 | 4.57912 | 2.34286 |
| 25 | PI16 | 7.34659 | 7.66718 | 7.0258 | 11.65659 | 8.4884 | 0.0272 | 0.09437 | 6.49419 | 1.3235 |
| 26 | RIMS2 | 2.3712 | 0 | 0.98341 | 2.74637 | 3.57539 | 5.23936 | 3.91391 | 4.24409 | 4.12594 |
| 27 | ZIC4 | 0.03826 | 0 | 4.68234 | 5.86901 | 6.44213 | 5.4917 | 5.53747 | 3.0907 | 0.26039 |
| 28 | KRT34 | 8.21182 | 5.06895 | 7.43262 | 9.67557 | 10.54596 | 10.0278 | 9.98273 | 6.68513 | 6.96506 |
| 29 | CNIH3 | 5.83328 | 4.89892 | 4.15134 | 6.99122 | 7.71871 | 8.23783 | 7.02207 | 8.28367 | 8.31302 |
| 30 | TMEM119 | 8.5114 | 7.63364 | 7.03669 | 8.67041 | 9.20167 | 11.4985 | 9.43299 | 9.19068 | 9.56658 |
| 31 | GIPR | 2.63412 | 2.22586 | 1.98241 | 4.58542 | 4.73663 | 4.35247 | 4.01302 | 2.58485 | 2.65045 |
| 32 | HOXB3 | 5.00602 | 5.42296 | 4.37375 | 7.08621 | 6.95517 | 7.89095 | 4.36348 | 7.42759 | 4.93412 |
| 33 | HOXD-AS2 | 5.05181 | 3.06928 | 5.93574 | 6.12601 | 8.12102 | 6.84048 | 6.20213 | 6.27605 | 6.53606 |
| 34 | SLC6A17 | 3.63612 | 3.60086 | 2.98163 | 5.79627 | 6.31519 | 2.84045 | 5.43317 | 4.77252 | 4.58779 |
| 35 | GSTM5 | 2.37309 | 0.90725 | 2.30363 | 3.93676 | 3.56838 | 4.61762 | 4.20071 | 7.04451 | 6.00616 |
| 36 | APBB1IP | 6.40944 | 6.23893 | 3.78881 | 8.96128 | 7.40586 | 6.31886 | 8.08079 | 7.18497 | 8.59549 |
| 37 | MACROD2 | 2.04828 | 2.4907 | 1.56786 | 4.0297 | 3.46256 | 4.55309 | 3.10603 | 3.10566 | 3.77197 |
| 38 | CTSK | 7.90371 | 7.20272 | 7.93561 | 7.76858 | 8.99564 | 10.80491 | 8.84005 | 8.99787 | 9.41024 |
| 39 | BATF3 | 2.85873 | 2.7074 | 1.56708 | 3.25833 | 5.72903 | 2.03277 | 5.57078 | 5.63231 | 6.68891 |
| 40 | HOXD4 | 4.37232 | 2.22276 | 3.30366 | 5.77169 | 5.73045 | 4.9864 | 5.01787 | 4.41681 | 2.665 |
| 41 | KRTAP2-3 | 8.19654 | 6.29124 | 6.37372 | 8.33241 | 9.42885 | 9.51318 | 9.00713 | 7.22591 | 8.75438 |
| 42 | SGCA | 4.37309 | 4.42406 | 0.98286 | 4.02391 | 7.12902 | 3.35516 | 5.53862 | 4.77055 | 5.66074 |
| 43 | TRPV2 | 6.49493 | 7.01792 | 7.55886 | 8.8547 | 9.72462 | 7.72696 | 8.55421 | 8.93935 | 9.0872 |
| 44 | HOXB2 | 7.92226 | 7.9598 | 5.0258 | 9.66383 | 9.32604 | 8.54518 | 7.54182 | 9.08894 | 6.63562 |
| 45 | LBH | 10.1301 | 7.82965 | 8.08469 | 10.66024 | 10.74434 | 11.17303 | 9.39874 | 8.71471 | 10.66394 |
| 46 | EMX2 | 8.25647 | 8.19371 | 1.9821 | 9.02858 | 9.21303 | 9.93728 | 6.60756 | 7.89587 | 9.20718 |
| 47 | RAC2 | 6.60657 | 7.68696 | 7.96443 | 9.65357 | 8.76434 | 9.41062 | 8.80539 | 7.83271 | 7.79269 |
| 48 | LOC101928324 | 2.63536 | 3.71038 | 2.3039 | 3.84535 | 5.98526 | 2.35441 | 4.27935 | 4.30131 | 4.03193 |
| 49 | NCAM1 | 5.54361 | 5.65429 | 1.98189 | 6.42804 | 6.42181 | 7.25228 | 7.08014 | 7.90386 | 6.43446 |
| 50 | ZNF79 | 2.05042 | 3.70949 | 3.68245 | 4.89316 | 5.39751 | 4.49141 | 4.75355 | 3.77359 | 4.50223 |
| 51 | GAS1 | 7.81995 | 6.43822 | 4.78876 | 7.94717 | 8.77683 | 8.5686 | 6.43747 | 7.21816 | 8.29076 |
| 52 | MFGE8 | 11.40459 | 10.97307 | 11.34445 | 12.21238 | 13.34723 | 12.86264 | 12.43141 | 11.99845 | 11.42243 |
| 53 | TMEM86A | 3.75082 | 2.70833 | 3.30379 | 4.84556 | 4.39488 | 5.38916 | 4.281 | 5.30181 | 4.14329 |
| 54 | DOCK2 | 3.21996 | 2.90215 | 1.98208 | 4.33153 | 4.73236 | 4.03181 | 3.1114 | 3.68272 | 5.71088 |
| 55 | WDR66 | 3.63658 | 4.3595 | 4.22953 | 4.52145 | 5.84671 | 6.1823 | 6.30196 | 5.92174 | 5.24933 |
| 56 | ACP5 | 2.04958 | 2.48798 | 0.98329 | 2.93856 | 3.96198 | 3.49108 | 5.46169 | 4.36739 | 4.3218 |
| 57 | CRYBB2 | 5.51042 | 0 | 0 | 0 | 7.09906 | 0.02556 | 6.06703 | 6.55369 | 6.22303 |
| 58 | ZNF169 | 3.36931 | 2.71344 | 0 | 3.64205 | 4.34016 | 4.11736 | 2.91379 | 2.589 | 3.63559 |
| 59 | HIVEP3 | 7.06296 | 5.73194 | 5.83941 | 8.06542 | 7.97561 | 7.33711 | 7.68478 | 7.57501 | 7.06265 |
| 60 | CCDC85A | 6.89725 | 6.66748 | 5.53601 | 7.44383 | 7.79185 | 8.43411 | 6.22393 | 8.35682 | 4.9941 |
| 61 | NEURL1B | 8.12343 | 7.34187 | 7.63962 | 7.20244 | 10.45701 | 7.77498 | 9.21622 | 10.68787 | 10.25192 |
| 62 | THY1 | 11.08885 | 11.06384 | 9.91804 | 11.3296 | 12.21132 | 12.81855 | 11.87183 | 12.25971 | 11.67836 |
| 63 | COX7A1 | 9.19151 | 8.36655 | 8.8883 | 9.70721 | 10.54756 | 10.49802 | 10.31769 | 9.37748 | 10.08967 |
| 64 | SPRY4-IT1 | 6.00601 | 5.9216 | 6.48929 | 8.05524 | 7.49659 | 7.07768 | 6.85642 | 7.30715 | 6.7742 |
| 65 | LINC00565 | 4.29932 | 4.14784 | 4.68212 | 5.2945 | 5.54151 | 6.40752 | 5.0677 | 5.85937 | 5.79977 |
| 66 | PKP1 | 6.00568 | 5.29181 | 4.37376 | 6.10623 | 7.82856 | 5.16263 | 6.34315 | 6.63949 | 6.36809 |
| 67 | FBLN5 | 8.75929 | 8.14217 | 8.1107 | 10.09062 | 9.14712 | 9.93883 | 8.82292 | 8.82457 | 9.17625 |
| 68 | HSD3B7 | 6.84635 | 7.4841 | 6.92394 | 8.37884 | 8.63155 | 8.54118 | 8.15971 | 8.75874 | 8.00732 |
| 69 | CRISPLD2 | 7.57541 | 6.96518 | 7.22937 | 8.94161 | 8.09546 | 8.81799 | 7.89009 | 8.02919 | 6.47924 |
| 70 | PPFIBP2 | 5.51109 | 4.75719 | 5.02587 | 4.7432 | 6.9013 | 6.97557 | 7.45362 | 6.58906 | 6.70293 |
| 71 | VTRNA1-2 | 2.85722 | 2.90284 | 2.30404 | 3.39808 | 4.8761 | 3.35386 | 5.01301 | 4.68921 | 3.65161 |
| 72 | LINC01224 | 2.04696 | 0 | 0 | 2.5268 | 2.46353 | 2.0303 | 4.48989 | 4.55505 | 2.89391 |
| 73 | ZFP37 | 4.80492 | 3.07047 | 3.56675 | 5.63918 | 5.04127 | 5.202 | 4.7552 | 3.47091 | 4.72847 |
| 74 | ICAM2 | 2.04761 | 0.91256 | 1.56807 | 3.26392 | 3.04824 | 2.03071 | 4.68395 | 3.10915 | 4.20823 |
| 75 | PSG1 | 3.7502 | 0.90838 | 0 | 2.93761 | 3.77851 | 4.20174 | 4.91542 | 4.63701 | 5.7462 |
| 76 | EMX2OS | 7.01768 | 7.88226 | 0.98298 | 8.28908 | 8.81226 | 9.24278 | 6.02256 | 7.20141 | 8.11031 |
| 77 | TLE1 | 5.95872 | 6.42293 | 6.39088 | 7.83106 | 7.39071 | 7.32788 | 7.23378 | 7.06237 | 7.11578 |
| 78 | TPST2 | 7.65925 | 6.84148 | 7.91217 | 8.37875 | 8.71826 | 9.18061 | 8.79836 | 7.91642 | 9.38234 |
| 79 | COL18A1 | 9.01773 | 9.08141 | 8.33014 | 9.56509 | 10.06602 | 10.43647 | 9.4532 | 9.68246 | 9.00819 |
| 80 | AKR1E2 | 3.21955 | 3.22521 | 3.15214 | 4.33215 | 4.83018 | 3.93845 | 3.1105 | 4.42497 | 4.23213 |
| 81 | HNMT | 5.44369 | 5.35944 | 5.30353 | 6.55159 | 5.99743 | 7.02145 | 5.94677 | 4.72197 | 5.51541 |
| 82 | ZNF530 | 3.50912 | 3.07245 | 0 | 4.64021 | 4.33317 | 1.61677 | 5.15112 | 3.94739 | 5.02244 |
| 83 | MAF | 3.22076 | 2.48581 | 3.56684 | 3.84437 | 4.45599 | 4.67597 | 0.08258 | 2.09319 | 6.86164 |
| 84 | LYL1 | 2.37194 | 2.90273 | 3.30416 | 4.46347 | 3.96092 | 3.83903 | 3.5692 | 5.0314 | 4.57109 |
| 85 | DKK1 | 9.75589 | 8.95421 | 9.80478 | 11.73255 | 12.06192 | 11.64959 | 10.00747 | 10.75791 | 10.13159 |
| 86 | COL16A1 | 9.93608 | 10.11947 | 8.99544 | 10.2281 | 9.48471 | 9.49291 | 8.49934 | 8.57381 | 7.4056 |
| 87 | TRIM61 | 3.7506 | 3.70959 | 3.30386 | 4.39794 | 5.15904 | 4.61682 | 4.56935 | 4.90732 | 3.65896 |
| 88 | OSBPL3 | 7.51127 | 7.20293 | 6.97012 | 8.47511 | 7.80323 | 8.72335 | 7.97944 | 7.85982 | 8.01414 |
| 89 | MAMLD1 | 7.84629 | 7.18446 | 6.82691 | 7.90736 | 8.69316 | 8.71639 | 7.74481 | 7.7724 | 9.54813 |
| 90 | PITX2 | 6.77963 | 3.48418 | 0.98292 | 8.24006 | 8.55401 | 7.78812 | 6.50749 | 6.66219 | 5.51962 |
| 91 | SNTA1 | 7.9223 | 8.39494 | 7.39081 | 9.09568 | 9.11913 | 9.01643 | 9.24726 | 8.90633 | 8.75465 |
| 92 | C17orf107 | 3.22013 | 3.71027 | 3.68259 | 4.84644 | 4.68103 | 5.64567 | 2.69707 | 3.35885 | 3.14434 |
| 93 | SOX4 | 10.20171 | 10.99243 | 10.11841 | 11.22919 | 10.55799 | 11.00144 | 8.94542 | 9.13921 | 8.63107 |
| 94 | LIMCH1 | 5.60659 | 8.17997 | 8.04754 | 9.07802 | 8.7584 | 9.71635 | 4.87057 | 7.05194 | 7.51953 |
| 95 | CDA | 3.75211 | 3.35884 | 3.78887 | 6.91488 | 6.66427 | 6.6764 | 3.81538 | 3.35226 | 5.09903 |
| 96 | HOXC5 | 0.03797 | 0 | 0.98291 | 5.52323 | 6.21237 | 6.57087 | 0.0847 | 0.7868 | 5.09119 |
| 97 | LIMK1 | 7.89725 | 7.10865 | 6.30336 | 8.45565 | 8.1004 | 8.41271 | 7.78763 | 7.5873 | 6.79242 |
| 98 | ANO3 | 2.63505 | 0 | 2.98212 | 5.40035 | 4.3336 | 5.64527 | 2.43326 | 1.77532 | 3.33295 |
| 99 | SGCG | 4.04925 | 1.48905 | 3.56716 | 5.36678 | 5.16097 | 4.88919 | 2.11132 | 2.35796 | 2.92118 |
| 100 | GAS7 | 6.81984 | 4.89898 | 3.78878 | 8.30269 | 8.02735 | 8.20808 | 6.02217 | 5.3516 | 4.59809 |
| 101 | HOXC4 | 2.37308 | 0 | 3.30339 | 7.62299 | 7.49713 | 8.61416 | 0.09194 | 4.08834 | 6.57565 |
| 102 | MYADM | 12.04315 | 11.39579 | 11.22636 | 12.79113 | 12.92434 | 12.24996 | 12.48393 | 11.87713 | 12.02137 |
| 103 | MMP17 | 5.09546 | 4.75831 | 4.5666 | 5.29443 | 6.09839 | 6.14113 | 5.8407 | 5.55236 | 4.73698 |
| 104 | DUSP14 | 9.81352 | 9.66384 | 9.32793 | 10.5966 | 11.05548 | 10.25871 | 10.27171 | 10.37932 | 10.42551 |
| 105 | EN1 | 4.57554 | 5.25646 | 5.7363 | 8.58707 | 9.54127 | 9.01344 | 7.2343 | 5.95623 | 4.34998 |
| 106 | UBE2QL1 | 2.36916 | 0 | 0 | 4.33681 | 4.58555 | 4.27678 | 2.10493 | 0 | 1.30178 |
| 107 | ABLIM3 | 8.4782 | 8.48775 | 7.36511 | 9.15977 | 9.44572 | 9.11028 | 10.04172 | 9.19922 | 9.08777 |
| 108 | TSHZ2 | 4.37362 | 4.22133 | 5.15153 | 7.08693 | 6.59676 | 7.3902 | 4.43691 | 4.41139 | 1.32699 |
| 109 | GSN | 9.85612 | 10.10718 | 11.02581 | 11.10874 | 11.62195 | 11.62175 | 9.86556 | 10.35976 | 11.12409 |
| 110 | RCAN3 | 5.77957 | 5.70689 | 5.68196 | 6.81903 | 6.66196 | 6.81449 | 6.43647 | 6.9978 | 6.83756 |
| 111 | HAND2 | 2.37325 | 2.70698 | 4.15144 | 7.52251 | 7.18262 | 6.26161 | 0.08926 | 3.57444 | 4.66856 |
| 112 | ADGRE5 | 8.78669 | 9.14927 | 9.22697 | 10.06886 | 10.80147 | 8.81814 | 9.96736 | 10.93713 | 10.20792 |
| 113 | HOXB7 | 0.04129 | 0.9073 | 0.98289 | 7.93646 | 7.54002 | 8.86589 | 0.0922 | 6.30684 | 0.2825 |
| 114 | CNNM1 | 3.50897 | 2.90258 | 0 | 0.9419 | 3.33112 | 4.83841 | 3.91656 | 5.17589 | 5.12717 |
| 115 | CCDC89 | 2.0498 | 2.22454 | 3.5673 | 3.74641 | 3.57306 | 3.93845 | 4.4997 | 5.10605 | 4.49317 |
| 116 | HOXB5 | 0.04198 | 2.48585 | 0 | 9.71086 | 9.59998 | 8.71283 | 1.10642 | 7.44637 | 0.28761 |
| 117 | RPSAP52 | 6.49457 | 6.10891 | 5.44089 | 7.71118 | 6.59501 | 6.33709 | 7.19136 | 6.85558 | 7.22907 |
| 118 | COL12A1 | 10.51651 | 10.49609 | 10.42953 | 11.14516 | 9.38187 | 12.32322 | 9.4842 | 8.79421 | 9.53512 |
| 119 | C12orf75 | 12.90803 | 11.51823 | 11.87167 | 12.88768 | 13.23889 | 13.1761 | 12.94682 | 13.65221 | 13.74207 |
| 120 | KCCAT198 | 3.36932 | 1.49193 | 0.98369 | 1.94009 | 3.04626 | 3.83751 | 4.19016 | 4.37617 | 3.77183 |
| 121 | HOXA11-AS | 0.03386 | 0 | 1.9823 | 5.43392 | 5.30515 | 4.83786 | 1.69364 | 1.36555 | 0.22862 |
| 122 | SLC27A1 | 5.95868 | 6.85382 | 5.59616 | 6.12577 | 7.66185 | 6.89126 | 7.6459 | 7.18595 | 7.10203 |
| 123 | MGAT5B | 4.04843 | 2.225 | 1.56745 | 2.5233 | 2.1935 | 4.93747 | 4.49876 | 4.42666 | 4.40838 |
| 124 | LOC148709 | 2.85519 | 1.49213 | 2.56773 | 3.11087 | 3.68618 | 2.353 | 3.91056 | 4.43618 | 3.63448 |
| 125 | MAPKAPK3 | 7.76616 | 8.12793 | 7.65386 | 8.35786 | 8.64784 | 8.58803 | 8.75201 | 9.03842 | 9.12716 |
| 126 | LINC00839 | 5.57491 | 3.48415 | 5.02596 | 4.52097 | 4.77393 | 6.52474 | 6.57285 | 6.64088 | 6.97118 |
| 127 | HOXC6 | 3.75195 | 3.80667 | 4.93565 | 11.17236 | 11.30253 | 11.83961 | 2.69784 | 4.29101 | 8.55481 |
| 128 | KHDRBS3 | 7.78651 | 5.54277 | 7.26686 | 7.67734 | 8.02655 | 7.50915 | 8.3959 | 8.57798 | 8.35967 |
| 129 | ENO3 | 3.75111 | 4.14981 | 3.88879 | 4.02492 | 3.95714 | 5.20189 | 4.57075 | 5.44947 | 5.28602 |
| 130 | SULF1 | 10.34315 | 10.94746 | 11.29089 | 11.77017 | 10.24014 | 11.91843 | 9.35411 | 9.42988 | 9.76177 |
| 131 | LINC00842 | 5.37237 | 5.45648 | 5.06935 | 6.86921 | 5.54259 | 3.61809 | 2.43597 | 2.354 | 2.34346 |
| 132 | SERPINF1 | 6.37355 | 6.86559 | 7.02598 | 6.75638 | 6.07633 | 8.27145 | 5.75904 | 4.57381 | 6.18078 |
| 133 | BEX1 | 8.07985 | 6.16569 | 5.68184 | 5.39511 | 9.01613 | 4.20331 | 10.80183 | 9.2282 | 10.26808 |
| 134 | LOC641367 | 2.37151 | 3.2261 | 3.30435 | 3.10884 | 3.04254 | 4.20093 | 4.27652 | 4.73934 | 4.32027 |
| 135 | HOXC-AS1 | 0.03391 | 0 | 0 | 5.33421 | 5.85739 | 4.27887 | 0.07552 | 0 | 1.31615 |
| 136 | DGAT2 | 3.95655 | 4.65939 | 4.37468 | 5.96301 | 3.95926 | 1.6169 | 2.43408 | 3.86293 | 2.33748 |
| 137 | HOXC9 | 1.63422 | 2.90052 | 3.98149 | 10.06256 | 10.57395 | 10.42573 | 2.43454 | 0.7883 | 7.1184 |
| 138 | PCOLCE2 | 8.67028 | 9.45952 | 8.60715 | 10.26679 | 8.98797 | 6.82793 | 7.92936 | 7.61822 | 7.46901 |
| 139 | HOTAIR | 0.03701 | 0 | 0 | 5.77223 | 6.50421 | 5.732 | 0.08253 | 1.77296 | 0.25133 |
| 140 | FOXD2 | 3.0501 | 3.36223 | 3.8892 | 4.10893 | 4.33296 | 1.6168 | 4.19659 | 5.13973 | 4.78388 |
| 141 | BMP2 | 5.51075 | 6.02982 | 6.76316 | 7.88578 | 4.03656 | 2.61819 | 3.70028 | 2.93721 | 4.99054 |
| 142 | ZNF714 | 5.83275 | 6.04991 | 4.06891 | 6.02391 | 5.99701 | 5.16256 | 6.89494 | 6.79237 | 6.51412 |
| 143 | HOXA11 | 0.03992 | 0 | 2.78888 | 7.05638 | 6.43906 | 7.7873 | 2.69969 | 0.7861 | 1.32699 |
| 144 | PARD6A | 5.18018 | 4.4238 | 4.7366 | 3.10601 | 5.97833 | 4.55659 | 6.22054 | 6.03939 | 5.55015 |
| 145 | VEGFA | 9.63504 | 10.30629 | 10.50008 | 10.77415 | 8.49189 | 10.84003 | 8.68173 | 8.56076 | 9.58607 |
| 146 | HOXC-AS3 | 0.03536 | 0 | 0 | 4.74713 | 5.33538 | 6.35233 | 0.07881 | 0 | 0.23943 |
| 147 | COL1A1 | 16.63677 | 16.59483 | 16.97586 | 16.68771 | 15.10063 | 17.57401 | 16.21628 | 14.64367 | 15.80766 |
| 148 | HOXA10 | 2.37266 | 1.48922 | 5.78876 | 9.52659 | 9.61841 | 9.68258 | 4.75954 | 3.35285 | 1.32437 |
| 149 | COL1A2 | 15.68825 | 16.27549 | 15.70411 | 15.57491 | 14.84843 | 16.48346 | 14.9197 | 14.25143 | 15.00408 |
| 150 | HOXA10-AS | 2.05054 | 1.4887 | 5.02581 | 8.39982 | 8.40273 | 9.04993 | 3.81597 | 1.77403 | 1.32567 |
| 151 | HOXC11 | 0.03915 | 0 | 0 | 6.67944 | 7.43384 | 6.07686 | 1.10715 | 0 | 1.32665 |
| 152 | MEI1 | 4.63259 | 3.81213 | 4.15266 | 4.69586 | 3.96232 | 2.61695 | 2.69512 | 3.09909 | 2.91587 |
| 153 | NRG4 | 4.37062 | 4.60595 | 4.30446 | 4.39907 | 3.77953 | 4.2794 | 2.69632 | 3.23457 | 3.65311 |
| 154 | VLDLR | 6.55903 | 6.45485 | 6.76305 | 6.32879 | 4.95427 | 7.01023 | 5.43671 | 5.15995 | 5.86982 |
| 155 | KCNIP4 | 4.29512 | 3.81357 | 3.88979 | 2.26101 | 3.96426 | 4.27839 | 2.69375 | 2.77871 | 3.13229 |
| 156 | STAC | 5.09508 | 5.80899 | 5.91276 | 4.52178 | 4.95657 | 6.09841 | 3.81471 | 2.09193 | 4.98491 |
| 157 | PELI1 | 6.52695 | 6.16694 | 6.959 | 6.89091 | 4.77363 | 6.31847 | 5.47225 | 5.41189 | 5.59462 |
| 158 | TMEM189 | 8.59491 | 9.06158 | 9.40563 | 9.25099 | 8.11428 | 8.43433 | 8.31516 | 7.85404 | 7.93474 |
| 159 | SYT15 | 5.72362 | 6.45551 | 6.78918 | 6.86786 | 4.62197 | 5.35503 | 5.0214 | 4.85566 | 4.25503 |
| 160 | HOXB-AS3 | 0.04007 | 0 | 0 | 6.31201 | 7.15496 | 8.02114 | 1.69656 | 0 | 0.27358 |
| 161 | NPR3 | 9.05752 | 10.02977 | 9.20545 | 9.46328 | 9.39782 | 7.25268 | 8.08714 | 7.43882 | 8.06418 |
| 162 | HOXA4 | 0.03868 | 0 | 3.56658 | 6.14754 | 7.24741 | 6.05462 | 0.08631 | 0 | 0.26346 |
| 163 | COL5A2 | 11.71909 | 11.86546 | 12.33208 | 11.30854 | 9.9403 | 12.24965 | 11.30427 | 10.07692 | 10.97505 |
| 164 | GLIS3 | 8.46958 | 8.86814 | 7.78883 | 8.28468 | 7.03636 | 7.98758 | 7.02234 | 6.50803 | 7.26814 |
| 165 | SCG2 | 7.81961 | 7.08955 | 7.78896 | 7.73722 | 5.74922 | 6.78817 | 5.63892 | 3.85393 | 5.80836 |
| 166 | LINC00673 | 4.75049 | 5.48852 | 5.40853 | 5.1478 | 1.86965 | 5.07651 | 4.35989 | 3.09181 | 3.80023 |
| 167 | FKBP11 | 9.23924 | 9.8299 | 9.47129 | 9.2258 | 8.98272 | 8.17304 | 8.63518 | 8.18436 | 8.32716 |
| 168 | DACT1 | 8.78643 | 8.89945 | 8.31688 | 8.76888 | 7.18838 | 7.57258 | 6.18173 | 5.28981 | 6.10502 |
| 169 | CLUHP3 | 4.57168 | 4.76449 | 4.23055 | 3.93944 | 3.7803 | 3.83891 | 3.27992 | 3.68504 | 0.23357 |
| 170 | COL3A1 | 13.54522 | 13.58182 | 14.20428 | 12.81084 | 11.65119 | 13.94697 | 12.11324 | 10.81137 | 11.58215 |
| 171 | DDX26B | 5.50923 | 4.94665 | 5.5969 | 5.4618 | 3.03781 | 4.42492 | 4.11292 | 4.35691 | 4.14828 |
| 172 | CCDC120 | 5.25899 | 4.94742 | 5.30421 | 3.74457 | 4.45554 | 4.73265 | 4.35965 | 4.58154 | 2.66435 |
| 173 | BAHCC1 | 7.61394 | 8.1617 | 8.18611 | 7.51342 | 6.63473 | 7.22319 | 6.62314 | 7.2185 | 6.96349 |
| 174 | PLXDC2 | 6.05151 | 7.84904 | 7.31261 | 6.73088 | 2.86693 | 7.13105 | 5.02211 | 2.57575 | 0.27968 |
| 175 | TMCC1-AS1 | 4.63346 | 4.99356 | 4.37485 | 3.74608 | 3.77935 | 3.61691 | 3.81016 | 3.68344 | 2.65785 |
| 176 | LOC101926940 | 4.69238 | 4.36415 | 4.98272 | 3.52335 | 2.87052 | 4.35345 | 3.43259 | 3.47588 | 3.79075 |
| 177 | PEG10 | 9.1837 | 9.25235 | 8.94724 | 7.62222 | 8.73681 | 7.73392 | 7.52499 | 8.95172 | 8.8006 |
| 178 | MYO15B | 4.74822 | 4.36538 | 4.07007 | 3.84636 | 3.45785 | 2.61707 | 3.80898 | 4.30516 | 1.91489 |
| 179 | FBXO34 | 4.37061 | 4.7131 | 4.23044 | 3.63914 | 3.04101 | 3.4914 | 3.69474 | 3.94797 | 4.41291 |
| 180 | EP300-AS1 | 3.9556 | 4.07519 | 4.23059 | 2.74564 | 3.45778 | 2.83942 | 4.27735 | 4.91359 | 3.13894 |
| 181 | C7orf31 | 4.50839 | 4.85939 | 4.37474 | 2.93766 | 3.33024 | 4.11929 | 3.11161 | 4.16965 | 4.78529 |
| 182 | DAPK1 | 4.63292 | 4.99462 | 3.98256 | 2.26013 | 3.96158 | 3.83886 | 2.11047 | 3.5854 | 3.49937 |
| 183 | HOXA7 | 0.0406 | 0 | 4.68193 | 6.95915 | 7.67627 | 8.07712 | 0.09065 | 0 | 0.27747 |
| 184 | HIPK1-AS1 | 4.21845 | 4.15314 | 4.88977 | 2.93831 | 3.45753 | 3.49123 | 3.8093 | 3.68499 | 3.91204 |
| 185 | MPZL3 | 4.13588 | 5.26546 | 4.50637 | 2.9385 | 3.87412 | 3.73196 | 3.10995 | 2.3599 | 1.91465 |
| 186 | GAREM | 6.695 | 6.83131 | 7.10069 | 6.04496 | 5.22464 | 5.78805 | 5.40011 | 5.25977 | 5.43439 |
| 187 | COL14A1 | 4.90845 | 5.99048 | 6.02658 | 2.25863 | 5.22791 | 4.73306 | 4.02039 | 3.3539 | 4.66206 |
| 188 | IGF1 | 3.21845 | 4.81595 | 5.19292 | 4.64236 | 1.87346 | 1.6162 | 0.07328 | 0 | 1.31305 |
| 189 | ARFGEF3 | 4.50739 | 4.95175 | 4.50638 | 2.52326 | 0.88002 | 4.61581 | 2.43248 | 2.35999 | 3.64939 |
| 190 | LOC100129550 | 5.95741 | 5.98914 | 5.65429 | 4.18428 | 4.25996 | 5.16212 | 4.43604 | 6.19908 | 4.73628 |
| 191 | HOXA-AS3 | 0.04065 | 0 | 4.68192 | 7.00285 | 7.72061 | 8.10421 | 0.09075 | 0 | 0.2778 |
| 192 | ANKRD62P1-PARP4P3 | 3.9554 | 4.6618 | 5.07053 | 3.5242 | 2.45651 | 3.73189 | 2.43236 | 2.77583 | 3.64888 |
| 193 | LOC344887 | 6.24082 | 6.79596 | 6.26716 | 5.18429 | 3.95411 | 5.78788 | 6.20094 | 5.0539 | 5.20382 |
| 194 | AOX1 | 8.36921 | 8.07417 | 9.00663 | 6.91302 | 7.68026 | 6.95234 | 7.95455 | 8.87318 | 7.56855 |
| 195 | PLA2G4C | 4.63393 | 4.54824 | 4.30428 | 1.93785 | 2.67713 | 4.03195 | 5.35522 | 2.35726 | 4.71957 |
| 196 | HOXA9 | 0.04121 | 0 | 3.30339 | 8.32031 | 8.04787 | 8.36392 | 0.09204 | 0 | 0.28196 |
| 197 | THSD4 | 8.3555 | 8.35886 | 8.42029 | 7.26676 | 7.34228 | 6.12096 | 7.77344 | 8.30266 | 6.49951 |
| 198 | C1S | 9.84473 | 10.89766 | 11.01279 | 9.7495 | 8.09499 | 9.54129 | 9.22154 | 8.31709 | 8.62165 |
| 199 | HOXA10-HOXA9 | 0.04121 | 0 | 3.30339 | 8.32031 | 8.05283 | 8.36392 | 0.09204 | 0 | 0.28197 |
| 200 | TNFAIP8 | 8.52348 | 8.55033 | 9.36736 | 7.95264 | 7.15174 | 7.20339 | 8.01019 | 7.0612 | 8.24994 |
| 201 | LOC100499484-C9ORF174 | 3.85532 | 4.49286 | 4.37536 | 1.52484 | 3.57554 | 2.83898 | 1.69299 | 4.03175 | 3.90503 |
| 202 | BCL2L11 | 4.9565 | 5.26135 | 5.86513 | 3.93752 | 3.03859 | 4.55576 | 2.92066 | 4.02086 | 3.66179 |
| 203 | C1R | 10.53979 | 11.18733 | 11.37589 | 9.96913 | 8.0609 | 10.09554 | 8.38249 | 8.07463 | 9.62856 |
| 204 | SMAD1-AS1 | 5.17775 | 5.11481 | 4.15222 | 2.74482 | 3.77833 | 3.03247 | 3.8111 | 4.53314 | 3.65658 |
| 205 | SUSD5 | 7.51916 | 7.99263 | 7.87032 | 5.02316 | 6.84421 | 6.05577 | 6.36311 | 6.43973 | 7.2178 |
| 206 | GSAP | 5.6352 | 5.99002 | 5.47384 | 5.1073 | 3.18931 | 2.61809 | 6.1111 | 3.46903 | 4.15103 |
| 207 | FAM169A | 5.13503 | 4.36503 | 4.30457 | 1.93836 | 2.87134 | 3.6167 | 3.80936 | 1.36445 | 4.56999 |
| 208 | HOXC10 | 0.04213 | 0 | 0 | 9.52088 | 10.09677 | 10.21575 | 0.09411 | 0 | 2.33976 |
| 209 | DOK3 | 4.13631 | 4.813 | 5.37528 | 1.93823 | 3.57279 | 3.20201 | 3.43209 | 3.36088 | 3.5015 |
| 210 | CHRNA5 | 4.85334 | 4.15632 | 4.07061 | 0 | 1.46085 | 3.73106 | 1.69205 | 2.95088 | 3.48903 |
| 211 | EFEMP1 | 11.09909 | 12.56722 | 12.06556 | 10.75426 | 8.11407 | 10.30498 | 8.6432 | 9.20509 | 10.49614 |
| 212 | IL26 | 5.8046 | 5.99082 | 6.23022 | 0 | 4.87015 | 2.35488 | 2.92145 | 2.0923 | 4.42767 |
| 213 | CADPS2 | 6.15958 | 6.34374 | 6.99307 | 4.39545 | 3.32598 | 4.28127 | 6.62098 | 5.57659 | 5.55446 |
| 214 | SH3RF2 | 6.819 | 7.00974 | 5.65409 | 1.93713 | 5.2251 | 2.84067 | 5.28427 | 6.47104 | 5.83609 |
| 215 | ADAMTS3 | 5.2573 | 5.07622 | 5.19229 | 3.10801 | 0.87925 | 2.61731 | 2.69641 | 2.09571 | 3.79006 |
| 216 | ITGB4 | 3.50609 | 4.08036 | 4.56878 | 2.26245 | 0 | 1.02919 | 3.56286 | 2.95512 | 2.9013 |
| 217 | SOBP | 8.01161 | 8.10912 | 9.12625 | 6.00187 | 6.24151 | 1.61695 | 6.60714 | 6.58722 | 5.15755 |
| 218 | PKNOX2 | 4.29477 | 4.23027 | 5.50729 | 1.93951 | 1.46066 | 0.02092 | 1.10266 | 1.77966 | 2.90921 |
| 219 | CRISPLD1 | 5.43919 | 4.42995 | 5.23105 | 1.93858 | 0 | 0.022 | 3.69346 | 2.58276 | 1.31683 |

**Table S2 Two-way ANOVA with Tukey post hoc test for HOXA9.** n=5.

a. ANOVA - HOXA9

|  | **Sum of Squares** | **df** | **Mean Square** | ***F*** | ***p*** |
| --- | --- | --- | --- | --- | --- |
| Scar type | 10.811 | 2 | 5.405 | 1.7749 | 0.1910 |
| Tension | 39.125 | 1 | 39.125 | 12.8471 | **0.0015**** |
| Scar type ✻ Tension | 5.739 | 2 | 2.869 | 0.9422 | 0.4037 |
| Residuals | 73.091 | 24 | 3.045 |  |  |

b. Post Hoc Comparisons - Scar type ✻ Tension (planned comparisons)

| **Comparison** | | | |  |  |  |  |  |
| --- | --- | --- | --- | --- | --- | --- | --- | --- |
| **Scar type** | **Tension** | **Scar type** | **Tension** | **Mean Difference** | **SE** | **df** | **t** | ***p*** |
| N | + | N | - | 3.27000 | 1.104 | 24.00 | 2.96273 | **0.0068**** |
| H | + | H | - | 2.43800 | 1.104 | 24.00 | 2.20891 | **0.0370*** |
| K | + | K | - | 1.14400 | 1.104 | 24.00 | 1.03650 | 0.3103 |

**Table S3 Two-way ANOVA with Tukey post hoc test for HOXC10.** n=5.

 a. ANOVA – HOXC10

|  | **Sum of Squares** | **df** | **Mean Square** | ***F*** | ***p*** |
| --- | --- | --- | --- | --- | --- |
| Scar type | 8.149 | 2 | 4.074 | 1.9331 | 0.1666 |
| Tension | 16.621 | 1 | 16.621 | 7.8860 | **0.0097**** |
| Scar type ✻ Tension | 3.144 | 2 | 1.572 | 0.7457 | 0.4851 |
| Residuals | 50.584 | 24 | 2.018 |  |  |

b. Post Hoc Comparisons - Scar type ✻ Tension (planned comparisons)

| **Comparison** | | | |  |  |  |  |  |
| --- | --- | --- | --- | --- | --- | --- | --- | --- |
| **Scar type** | **Tension** | **Scar type** | **Tension** | **Mean Difference** | **SE** | **df** | **t** | ***p*** |
| N | + | N | - | 2.02600 | 0.9182 | 24.00 | 2.20653 | **0.0372*** |
| H | + | H | - | 1.86200 | 0.9182 | 24.00 | 2.02791 | 0.0538 |
| K | + | K | - | 0.57800 | 0.9182 | 24.00 | 0.79535 | 0.4342 |

**Table S4 Two-way ANOVA with Tukey post hoc test for COL1A1.** n=4,4,3 for normal, hypertrophic, and keloid fibroblasts.

 a. ANOVA – COL1A1

|  | **Sum of Squares** | **df** | **Mean Square** | ***F*** | ***p*** |
| --- | --- | --- | --- | --- | --- |
| Scar type | 0.9094 | 2 | 0.4547 | 0.4734 | 0.6313 |
| Tension | 19.8634 | 1 | 19.8634 | 20.6793 | **0.0003***** |
| Scar type ✻ Tension | 2.0880 | 2 | 1.0440 | 1.0869 | 0.3609 |
| Residuals | 15.3687 | 16 | 0.9605 |  |  |

b. Post Hoc Comparisons - Scar type ✻ Tension (planned comparisons)

| **Comparison** | | | |  |  |  |  |  |
| --- | --- | --- | --- | --- | --- | --- | --- | --- |
| **Scar type** | **Tension** | **Scar type** | **Tension** | **Mean Difference** | **SE** | **df** | **t** | ***p*** |
| N | + | N | - | 2.7350 | 0.6930 | 16.00 | 3.9465 | **0.0012**** |
| H | + | H | - | 1.8088 | 0.6930 | 16.00 | 2.6100 | **0.0190*** |
| K | + | K | - | 1.2100 | 0.8002 | 16.00 | 1.5121 | 0.1500 |

**Table S5** Profiles of fibroblasts from patient groups. M and F stands for male and female.

|  | Sex | Age | Ethnicity | Body site | Scar etiology |
| --- | --- | --- | --- | --- | --- |
| Normal Skin | M | 5 | Asian | Preauricular area | Skin graft donor |
|  | F | 15 | Asian | Thigh area | Skin graft donor |
|  | M | 30 | Asian | Retro auricular area | Skin graft donor |
| Hypertrophic scar | F | 10 | Asian | Thigh area | Surgery scar |
|  | F | 21 | Asian | Thigh area | Surgery scar |
|  | M | 30 | Asian | Upper lip | Trauma |
| keloid | F | 31 | Asian | Auricular helix | Ear ring piercing |
|  | M | 16 | Asian | Mastoid area | Surgery scar |
|  | F | 25 | Asian | Ear lobe | Ear ring piercing |

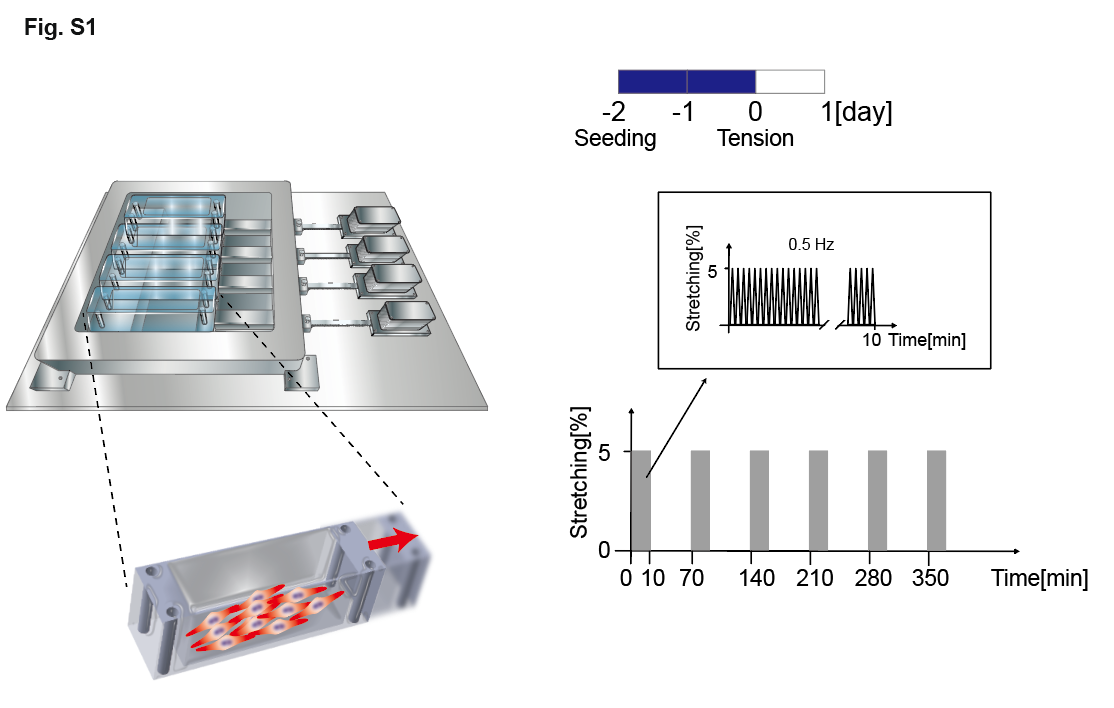

**Fig. S1** **In vitro tensile stimulation apparatus and the experimental condition**. Intermittent uni axial stimulations (5% strain, 0.5Hz) were used where each cycle consisted of a 10-minute stretching followed by a 60-minute resting period. Total 6 cycles were applied to each experiment.
